# Complementary coding of behaviors in striatal pathways supports a dual selection-suppression function

**DOI:** 10.1101/2022.06.20.496781

**Authors:** Christophe Varin, Amandine Cornil, Delphine Houtteman, Patricia Bonnavion, Alban de Kerchove d’Exaerde

## Abstract

The basal ganglia are known to control actions and modulate movements. Neuronal activity in the two efferent pathways of the dorsal striatum, a major input to the basal ganglia, is critical for appropriate behavioral control. Previous evidence has led to divergent conclusions on the respective engagement of both pathways during actions. We used calcium imaging to evaluate how neurons in the direct and indirect pathways in the dorsal striatum encode behaviors during self-paced spontaneous explorations in an open field. We observed that the two striatal pathways exhibit distinct tuning properties during spontaneous behaviors. We applied supervised learning algorithms and found that direct pathway neurons encode behaviors through their activation, whereas indirect pathway neurons exhibit behavior-specific silencing. These properties remain stable for weeks. Our findings highlight a complementary encoding of behaviors in the two striatal pathways that supports an updated model, reconciling previous conflicting conclusions on motor encoding in the striatum.

## INTRODUCTION

The basal ganglia are certainly well known to control both goal-directed behaviors and natural, self-paced behaviors. The proper initiation and execution of these behaviors relies heavily on appropriate functioning within the basal ganglia, as basal ganglia dysfunction is at the core of various disorders, including Parkinson’s disease, autism spectrum disorders, and schizophrenia^1^. The striatum, which is the main entry nucleus of the basal ganglia, consists of two types of striatal projection neurons (SPNs) that differ based on their expression of either dopamine D1 or D2 receptors and their respective direct or indirect projections to the output nuclei of the basal ganglia (dSPNs or iSPNs). This functional organization provides differential control of basal ganglia outputs by dSPNs or iSPNs, that leads to net activating or inhibiting effects of thalamocortical circuits, respectively^2, 3^. This dichotomy between prokinetic dSPNs and antikinetic iSPNs has been documented using loss-of-function or gain-of-function experiments^4–10^, bolstering the traditional go/no-go description of striatal functioning^2, 3^. This view has been challenged by correlative descriptions of striatal activity based on recordings of both types of SPNs, which demonstrated a coactivation of the dSPN and iSPN pathways during locomotion and more generally during actions^11–16^. These results indicate that concerted and cooperative activity between both striatal pathways is needed for proper action initiation and execution.

As a result, two antagonistic models of striatal functioning have been developed that explain how neuronal activity is organized in the striatum. The selection-suppression model^17^ postulates that proper action execution relies on the concurrent activation of a small discrete subpopulation of dSPNs that encode the ongoing action and the widespread activation of a large number of iSPNs that inhibit all other actions, with the iSPNs associated with the ongoing action remaining silent. This model predicts that dSPNs are more selective for actions than iSPNs. However, recent investigations highlighted that both pathways encode behaviors with similar properties and dynamics^13–16^. Consequently, a cooperative selection model was proposed^13^ in which dSPNs and iSPNs coordinate their activities to select the proper action, with subsets of dSPNs and iSPNs displaying the same targeted activation patterns toward actions. Although this model considers the coactivation of small ensembles of dSPNs and iSPNs, this model likely fails to take into account the functional opposition between dSPNs and iSPNs^4–10^. In summary, these models predict different patterns of neuronal activation in response to various behaviors, particularly the activity of iSPNs. Therefore, additional investigations are needed to clarify the function of the two SPN pathways as well as their relative organization into functional subpopulations for behavior encoding.

Here, we studied the behavior-encoding properties of dSPNs and iSPNs in the dorsal striatum using one-photon microendoscopy in mice that freely explored an open field and thus expressed a large behavioral repertoire at their own pace. We observed that the behavior-encoding properties of dSPNs and iSPNs differ in a way that challenges the above models. Furthermore, we used support vector machine classifiers to precisely analyze the neural code of dSPNs and iSPNs and their activation patterns during behaviors. We found that, despite their differences in encoding properties, both populations contain the same amount of information to reliably infer behaviors. Moreover, we classified neurons as activated or silent during behaviors to evaluate the predictions of the selection-suppression and cooperative selection models, and we found that neural codes are organized differently in dSPNs and iSPNs. Similar to previous observations in different brain systems, the most important behavior-encoding feature of dSPNs is their specific activation during some behaviors. Remarkably, the most important behavior-encoding feature of iSPNs is their consistent silencing during specific behaviors. Our findings are reinforced by observations that these properties remain stable for weeks. These results provide the first correlative evidence that dSPNs and iSPNs have distinct encoding properties, supporting an updated model for motor encoding among SPNs in the dorsal striatum that relies on the congruent activation of dSPNs, which encode multiple accessible behaviors in a given context to promote these behaviors, and iSPNs, which encode for and inhibit competing behaviors. As a result, the coactivation of specific subsets of dSPNs and iSPNs would result in the selection and execution of only one motor program. This updated model bridges the gap between various interpretations of experimental observations that promoted antagonistic models on striatal functional organization.

## RESULTS

### Behavior encoding properties differ between dSPNs and iSPNs

To investigate the behavior-encoding properties in both subpopulations in the dorsal striatum, we tracked and reconstructed the behavior of mice freely exploring an open field and simultaneously recorded the neuronal activity of either dSPNs or iSPNs using microendoscopic one-photon imaging of GCaMP6s (Fig. 1a). Mouse self-paced behaviors were identified and labeled using a combination of deep learning tools and clustering methods to generate a predictive model (Fig. 1b, Extended Data Figs. 1 and 2a-c). The behavior distributions of the experimental groups were similar. The calcium activity was extracted from simultaneously recorded microendoscopic images using the CaImAn pipeline^18, 19^, and the reconstructed temporal traces of calcium activity were deconvolved using MLspike^20^ (Fig. 1c-f, Extended Data Figs. 1 and 2d-k). On average, 179 ± 18 dSPNs and 216 ± 16 iSPNs were identified in each recording session (Fig. 1f). First, we observed that the average population activity of the dSPNs was higher than that of the iSPNs (Fig. 1f). Then, the population activity was decomposed according to the identified behaviors, which revealed that the average population activity was consistently higher for dSPNs than for iSPNs during many behaviors, including straight locomotion, locomotion with right and left turns, remaining still with right and left turns, rearing, grooming, and locomotion sniffing, with the notable exception of immobility, during which the average population activity was significantly lower for dSPNs than for iSPNs (Fig. 1g). This result indicates a substantial difference in the behavioral tuning properties of dSPN and iSPN ensembles.

**Fig. 1:**
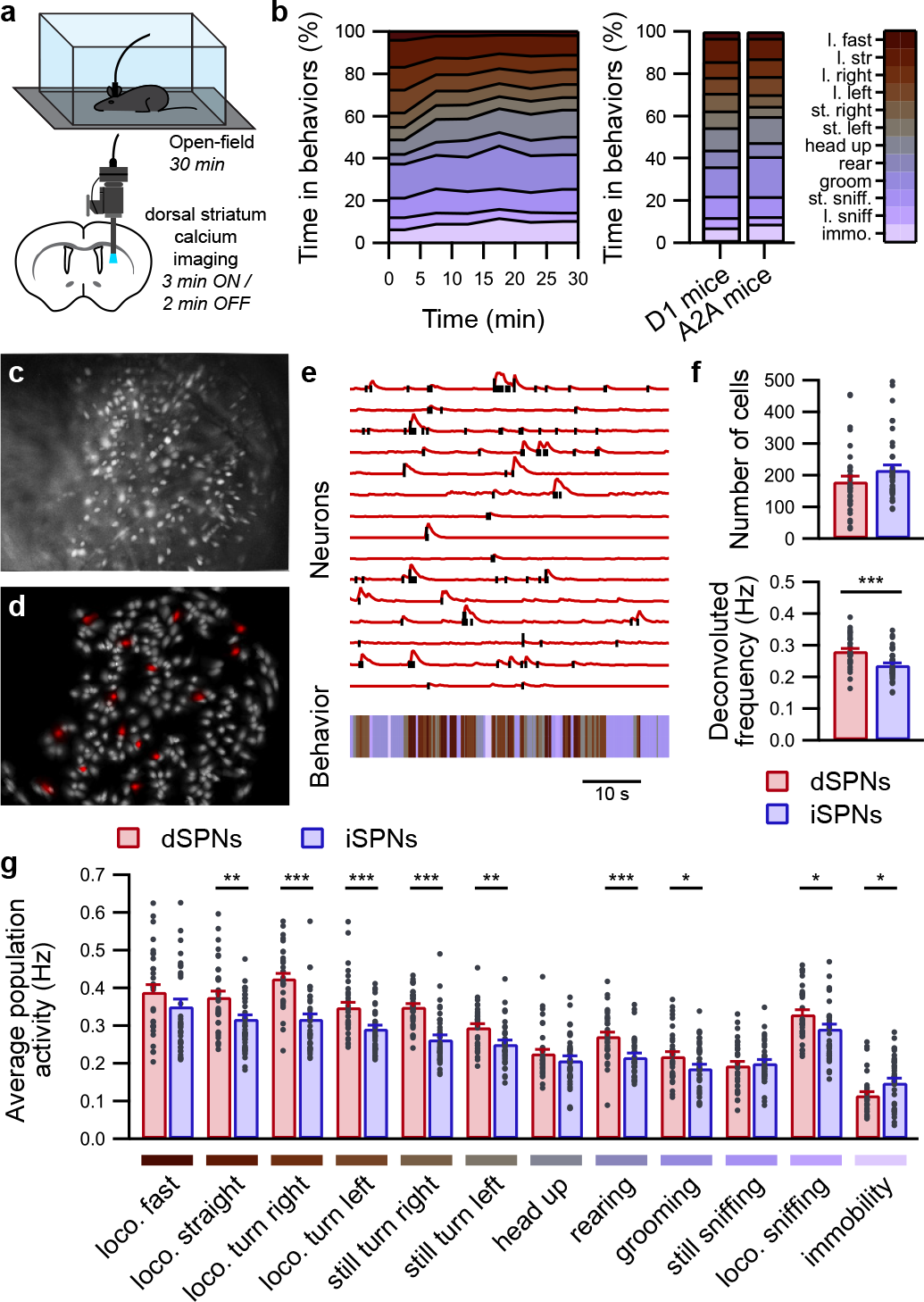
Simultaneous calcium imaging and behavioral identification in freely behaving mice. **a**, Mice expressing GCaMP6s in either dSPNs or iSPNs and equipped with a microendoscope freely exploring a well-known open field. **b**, Temporal evolution of identified behaviors during 30 min of open-field exploration for all recording sessions (left panel; 5 min long bins; n = 73 sessions in 17 mice) and average distribution of behaviors over 30 min (right panel) in mice expressing GCaMP6s in dSPNs (D1 mice; n = 33 sessions with 8 mice) or iSPNs (A2A mice; n = 40 sessions in 9 mice). Abbreviations: l., locomotion; st., still; immo., immobility. **c-d**, Representative image from one A2A mouse of the maximum fluorescence intensity projection of iSPNs labeled with GCaMP6s (c) and the corresponding isolated spatial components identified using CNMF-E (d). **e**, Representative fluorescence traces (top panel, red lines) and deconvolved calcium activity (top panel, black lines) from selected SPNs (red in d) aligned with detected behaviors (bottom panel). **f**, Quantification for each recording session of the identified neuron number (top panel) and deconvolved calcium activity (bottom panel) for dSPNs (red; n = 33 sessions in 8 mice) and iSPNs (blue; n = 40 sessions in 9 mice) (dSPNs vs. iSPNs: *** p < 0.001). **g**, Average population activity is significantly higher in dSPNs than in iSPNs during many behaviors and significantly lower during immobility (dSPNs vs. iSPNs: * p < 0.05, ** p < 0.01, *** p < 0.001). Abbreviation: loco., locomotion.

To better characterize this potential difference among the SPN subpopulations, we first evaluated whether SPN activation was consistent during 30 min of open-field exploration. For each behavior, we computed the neuronal activation, which was calculated as the average frequency, during the first and the second halves of each recording, and we evaluated the similarity between these two neuronal activation maps (Fig. 2a). We observed that neuronal activation similarity was higher for dSPNs than for iSPNs for all identified behaviors except grooming, still sniffing, and immobility (Fig. 2b, Extended Data Fig. 3a,b). This result suggests that, for each behavior, the same dSPNs are more consistently activated, whereas iSPN activation is inconsistent. Conversely, this observation between the first and second halves of the recordings could result from the fact that both subpopulations are differentially affected by internal drives that accumulate or dissipate during open-field exploration, such as stress^21^, novelty^22, 23^, or tiredness^24^. To account for this temporal factor, we computed the same similarity metric by using all possible partitions of time into two 15-min sets (e.g., 30 min segmented into 5 min long slices). Regardless of the time partition, we observed the same difference in neuronal activation similarity between dSPNs and iSPNs (Extended Data Fig. 3c). Furthermore, the neuronal activation similarity was computed by comparing odd and even frames and by comparing one episode out of two to complementary episodes, yielding the same observations as described above (Extended Data Fig. 3d,e). Moreover, as an additional control to verify the reliability of the similarity measure, we disturbed SPN signaling by artificially increasing dopamine release through acute injection of amphetamine. The neuronal activation similarity of both dSPNs and iSPNs was strongly alleviated after amphetamine administration (Extended Data Fig. 4a,b). All the above results demonstrate that the difference in neuronal activation similarity is a substantiated time-invariant property of behavior encoding in dSPNs and iSPNs; for each behavior, the same dSPNs are more consistently activated, whereas iSPN activation displays either a milder specificity toward behaviors or is more variable.

**Fig. 2:**
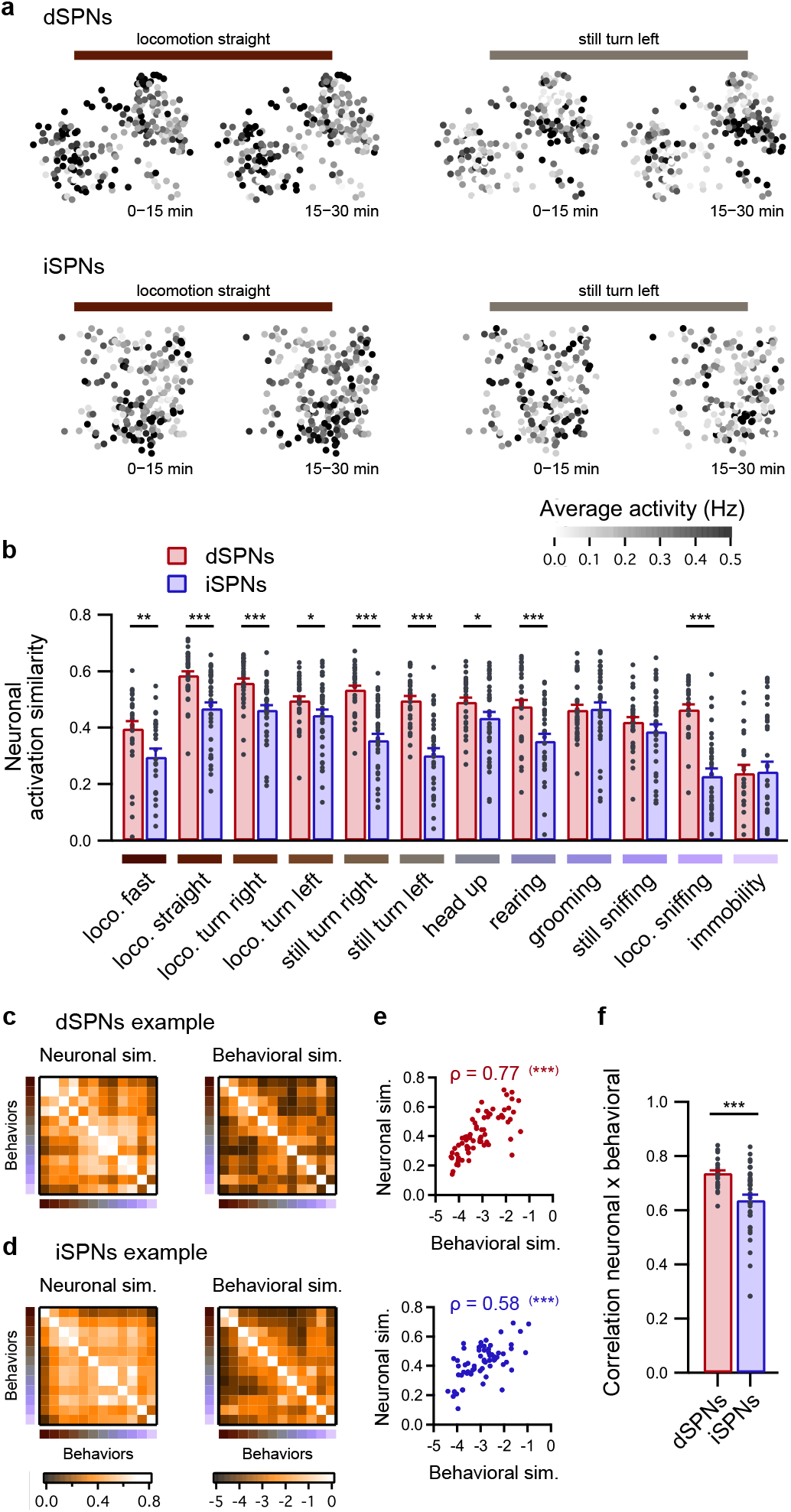
Dynamical properties of behavior encoding differ between dSPNs and iSPNs. **a**, Representative neuronal activation maps of dSPNs (top panels) and iSPNs (bottom panels) from one session, illustrating the averaged activation of neurons in the first half (0-15 min) and second half (15-30 min) of the recording session for two behaviors (locomotion straight, left; still turn left, right). Note that the neuronal activation appears highly similar during the first and second halves of the recording in dSPNs during the two behaviors and less similar in iSPNs during the same behavior. **b**, Neuronal activation similarity between the first and second halves of the open-field exploration was higher in dSPNs (red; n = 33 sessions in 8 mice) than in iSPNs (blue; n = 40 sessions in 9 mice) for all behaviors except grooming, still sniffing, and immobility (dSPNs vs. iSPNs: * p < 0.05, ** p < 0.01, *** p < 0.001). **c-d**, Examples of matrices of pairwise neuronal activation similarity between behaviors (left panels) and matrices of pairwise behavioral similarity between behaviors (right panels) in one session from one representative dSPNs recording (**c**) and one representative iSPNs recording (**d**). **e**, For pairs of behaviors, the neuronal activation similarity and behavioral similarity are significantly correlated in both dSPNs and iSPNs, as illustrated by the examples displayed in c-d (Spearman correlation: *** p < 0.001). **f**, Average correlation coefficient (Spearman correlation) between pairwise behavioral and neuronal similarities is higher for dSPNs (red; n = 33 sessions in 8 mice) than iSPNs (blue; n = 40 sessions in 9 mice) (dSPNs vs. iSPNs: *** p < 0.001).

Moreover, a comparison of pairs of behaviors revealed that in the dorsal striatum, similar behaviors are encoded by highly similar neuronal ensembles, whereas dissimilar behaviors are encoded by mildly overlapping neuron groups. Furthermore, this encoding property is equivalent for both SPN subtypes^14^. For each pair of behaviors, we compared the similarity between the average neuronal activity (neuronal activation similarity) to the similarity between behaviors (behavioral similarity). The latter quantifies the similarity in movement trajectories and body shape between each pair of behaviors. We observed a strong positive correlation between pairwise neuronal activation similarity and pairwise behavioral similarity for both dSPNs and iSPNs (Fig. 2a,c-e). However, when the correlation coefficients of pairwise neuronal and behavioral similarities were compared, we observed a significantly higher correlation for dSPNs than for iSPNs (Fig. 2f). This result indicates that the behavioral space representation differs between the two populations. This result contradicts that of a previous report^14^. To understand this result, we first verified that this difference was not due to the use of a different neuronal similarity measure (Extended Data Fig. 5a). Another key difference is the duration over which animals explored the open field: the animals explored the field for 10-15 min in Klaus, Martins et al., 2017^14^, whereas our experiments lasted 30 min. Thus, we calculated the correlation coefficients between the neuronal and behavioral similarities for different recording lengths. No difference was detected between dSPNs and iSPNs for durations of up to 10 min (Extended Data Fig. 5b-d). Therefore, extended recordings may be required to collect more samples of neuronal activity during a larger variety of internal or external contexts in order to properly uncover differences in neuronal activation variability for episodes of different behaviors. Overall, these results demonstrate that the coupling between behavior similarity and neuronal similarity is tighter among dSPNs than among iSPNs.

The above observations demonstrate fundamental differences between dSPNs and iSPNs in terms of their dynamical behavior-encoding properties. Moreover, this new set of evidence is inconsistent with existing models of striatal organization. In particular, these models do not account for the existence of different neuronal activation patterns during different episodes of the same behavior. Thus, our results call for a deeper analysis of the neural codes and properties of dSPNs and iSPNs that may convey different behavior-relevant information.

### Behaviors can be reliably decoded from either dSPNs or iSPNs

The first step in our analysis of the neural code in the dorsal striatum was to assess whether animal behaviors can be reconstructed based on instantaneous neuronal activity recorded from either dSPNs or iSPNs. We thus trained several support vector machine linear classifiers for each pair of behaviors and used the majority rule to combine the outputs of individual classifiers to predict animal behavior (one vs. one multiclass support vector machine) (Fig. 3a). The behavior reconstruction error was calculated for each prediction using the behavioral distance between the predicted behavior and the actual behavior. The instantaneous prediction of behaviors using either dSPN or iSPN activity performs significantly better than chance, which was estimated by decoding behaviors using classifiers trained on time-lagged data. Moreover, dSPNs and iSPNs have similar decoding accuracies and average reconstruction errors (Fig. 3b,c). The prediction accuracy associated with individual classifiers (separation between pairs of behaviors) is similar for dSPNs (81.8 ± 1.1%) and iSPNs (82.5 ± 0.8%) (p = 0.295). Strikingly, this result appears to contradict previous observations, which indicate differences in behavior encoding between dSPNs and iSPNs. A lower prediction accuracy would be expected when using iSPNs (because of their lower activation similarity) and a larger prediction error would be expected when using iSPNs (because of the lower coupling between pairwise neuronal and behavioral similarities). To confirm the relevance and robustness of our classification strategy, we evaluated whether the decoding accuracy is affected when SPN activity is strongly disturbed after amphetamine administration. We observed that the decoding performance was considerably reduced following amphetamine administration (Extended Data Fig. 4a, c-d). A closer inspection of the performance of individual classifiers revealed that the decoding accuracy was higher when distinguishing dissimilar behaviors, and lower when distinguishing similar behaviors (Fig. 3d). This property is the same for dSPNs and iSPNs. For example, for dSPNs, the separation between considerably different behaviors, such as locomotion turn right and immobility (behavior distance: 3.93 ± 0.06), is more accurate (accuracy: 90.2 ± 1.1%) than the separation between more similar behaviors, such as locomotion turn right and locomotion turn left (behavior distance: 1.79 ± 0.06; accuracy: 76.1 ± 1.3%). These results indicate that despite previous evidence on differences in dynamical behavior encoding, both striatal populations contain and encode the same amount of information in response to behaviors.

**Fig. 3:**
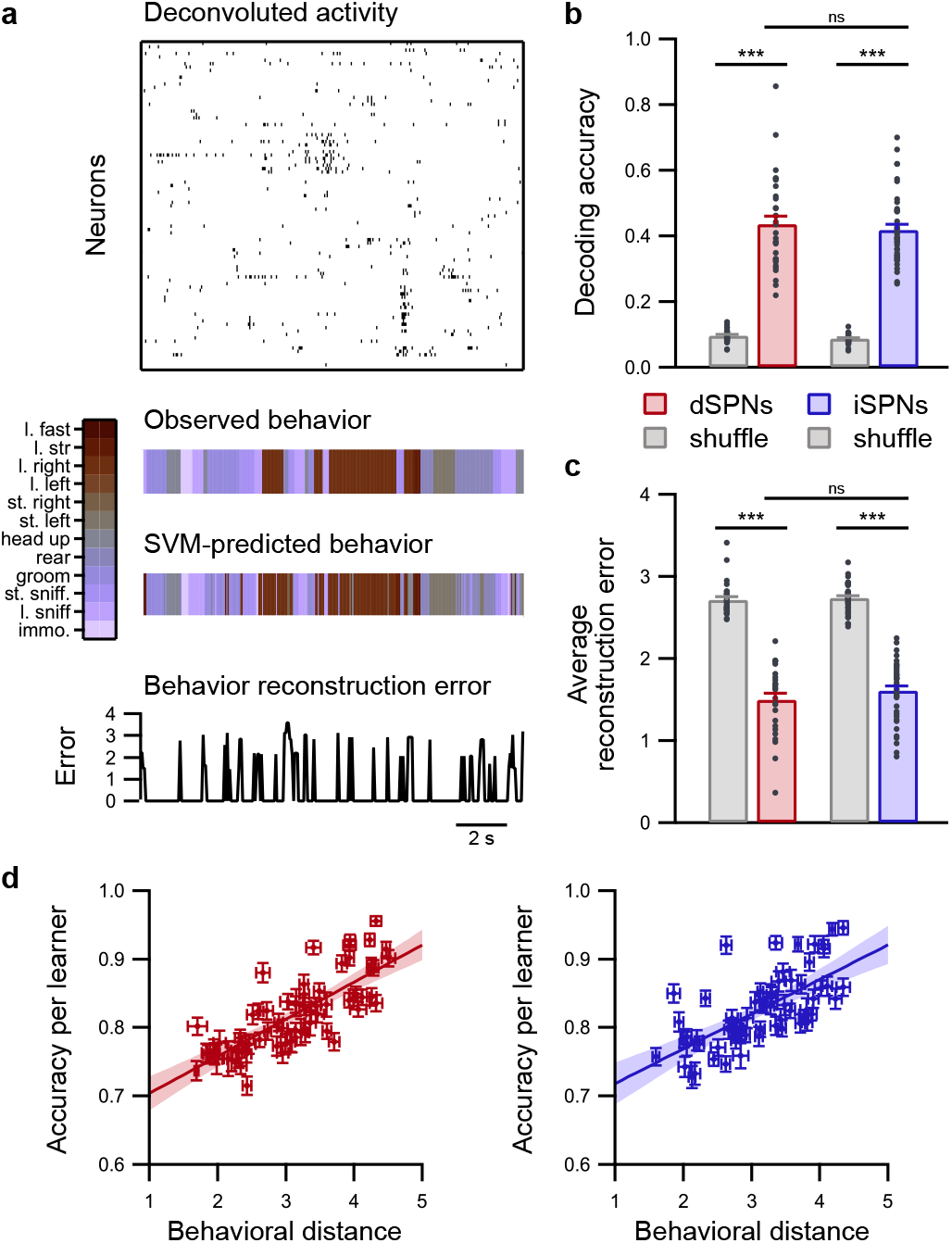
Predicting behaviors is equally efficient when using either dSPN or iSPN ensembles. **a**, Example of behavior decoding using support vector machines (SVM). Using deconvolved calcium signal (top panel), support vectors are trained to predict ongoing behaviors (middle panel). The prediction error (bottom panel) is calculated by using the behavioral distance between the observed and predicted behaviors. **b**, Behavior decoding accuracy was similar when dSPNs (red; n = 31 sessions in 8 mice) or iSPNs (red; n = 38 sessions in 9 mice) were used (dSPNs vs. iSPNs: ns p > 0.05). Gray, chance level when decoding from time-lagged data (dSPNs or iSPNs vs. shuffle: *** p < 0.01). **c**, Decoding error evaluated as the average behavioral error for the entire time series using dSPNs (red; n = 31 sessions in 8 mice) or iSPNs (red; n = 38 sessions in 9 mice) (dSPNs vs. iSPNs: ns p > 0.05). Gray, chance level when decoding from time-lagged data (dSPNs or iSPNs vs. shuffle: *** p < 0.01). **d**, Relationship between the behavioral distance and decoding accuracy for each individual SVM binary classifier using dSPNs (left panel) or iSPNs (right panel). The data for each classifier are presented as the mean (dots) ± SEM across sessions for both the behavioral distance and accuracy. Colored straight lines represent the average regression line, and shaded areas illustrate the 95% confidence interval.

### Neural code in dSPNs is biased toward activation

To better characterize the neural code in the dorsal striatum, we attempted to identify which features in the response properties of individual cells contribute most to behavior encoding. We first identified which neurons were significantly activated during specific behaviors and defined these neurons as behavior-active when the mutual information between the behavior time series and the recorded neuronal activity (behavior information) was statistically significant^25, 26^ (Fig. 4a,b, Extended Data Fig. 6a,b). Based on this criterion, the proportion of behavior-active cells is higher among dSPNs than among iSPNs (Fig. 4b, Extended Data Fig. 6c). According to this classification of whether individual neurons are behavior-active, we evaluated whether behaviors could be decoded based on the activity of behavior-active or non-behavior-active neurons. For dSPN recordings, the prediction was significantly more efficient when using behavior-active cells than when using non-behavior-active cells, whereas, for iSPN recordings, the prediction performance was similar when either behavior-active cells or non-behavior-active cells were used (Fig. 4c,d, Extended Data Fig. 6d,e). Moreover, the decoding performance was similar when non-behavior-active dSPNs or iSPNs were used, whereas the decoding performance was significantly higher when behavior-active dSPNs were used than when behavior-active iSPNs were used (Fig. 4c,d). This observation was maintained when the level of significance of behavior information for classifying neurons as behavior-active was changed (Extended Data Fig. 6f). Taken together, these findings demonstrate that the neural code for behaviors is biased toward behavior-active dSPNs, whereas this feature is more evenly distributed among iSPNs.

**Fig. 4:**
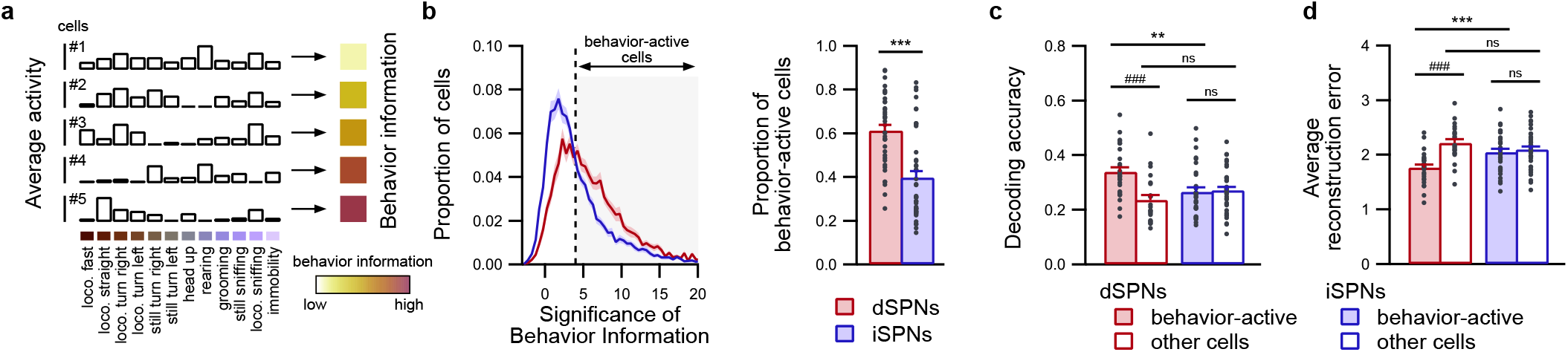
Relevant information for decoding behaviors lies primarily in excited dSPNs. **a**, Information content of neuronal activity in response to behaviors was derived based on the behavior information, which was calculated as the mutual information between the behavior time series and neuronal activity. As illustrated for 5 representative neurons, a higher level of behavior information reflects a stronger degree of tuning of neuronal activation toward behaviors. **b**, Cells are identified as behavior-active when the level of significance of behavior information exceeds 4 sigma of the shuffled distribution (gray area; left panel; threshold indicated by the black dashed vertical line). A larger proportion of behavior-active cells was identified among dSPNs (n = 29 sessions in 8 mice) than iSPNs (n = 37 sessions in 9 mice) (right panel; dSPNs vs. iSPNs: *** p < 0.001). **c-d**, Decoding accuracy (**c**) and average decoding error (**d**) when predicting behaviors using behavior-active neurons (plain bars) and non-behavior-active cells (unfilled bars). The decoding performance is better for behavior-active cells than for non-behavior-active cells in dSPNs recordings (red bars; n = 29 sessions in 8 mice), whereas it is similar in iSPNs recordings (blue bars; n = 37 sessions in 9 mice) ensembles. Moreover, the decoding performance is better for behavior-active dSPNs than for behavior-active iSPNs (dSPNs vs. iSPNs: ns p > 0.05, ** p < 0.01; behavior-active vs. other cells: ns p > 0.05, ^###^ p < 0.001).

### Neural code in iSPNs is biased toward silencing

According to the selection-suppression model^17^, indirect pathway SPNs are activated to suppress all motor programs except the one being executed. As a result, iSPNs responsible for suppressing a particular behavior are never active during this behavior. Thus, following this postulate, we investigated the relative contribution of neurons that remain silent during behaviors to the neural code. We thus identified SPNs that are consistently silent during episodes of each behavior by computing, for each neuron, how often this neuron is active during all episodes of this behavior (Fig. 5a). For each behavior, if this activation occurrence is sufficiently low (threshold of 2.5% of episodes; Fig. 5a), the neuron is labeled as behavior-silent for this behavior. Strikingly, the proportion of behavior-silent neurons was higher in iSPNs than in dSPNs for each behavior (Fig. 5b), with the notable exception of immobility, for which the proportion was similar in dSPNs and iSPNs. Then, we separated SPNs classified as behavior-silent or non-behavior-silent and evaluated the information content of these groups of cells to accurately separate the associated behavior from other behaviors (one behavior vs. rest decoding accuracy) (Fig. 5c). For dSPNs, the separation accuracy of all behaviors was similar when behavior-silent cells or non-behavior-silent cells were used, except fast locomotion and immobility. On the other hand, we observed that the separation accuracy was consistently higher for behavior-silent iSPNs than non-behavior-silent iSPNs during ambulatory behaviors (e.g., behaviors involving locomotion or right and left turns without locomotion), indicating that silencing is an important feature of the neural code in iSPNs during these behaviors. Conversely, this difference in separation accuracy between behavior-silent and non-behavior-silent iSPNs was not observed for static behaviors (e.g., head up, rearing, grooming, still sniffing, and immobility). This distinction between ambulatory and static behaviors could be because the detected static behaviors may contain additional substates that were not separated in our classification. Furthermore, this distinction may represent different modes of encoding in iSPNs during ambulatory and static behaviors, such as spatial mapping in the dorsal hippocampus, which is characterized by place cells that display their specific place fields mostly during locomotion^27, 28^. Additionally, we observed the same results for both dSPNs and iSPNs when we used an alternate cell classification, which was based on a threshold on the average activity during behaviors (Extended Data Fig. 7). In conclusion, these observations support a key distinctive feature of iSPNs for behavior encoding, namely, that iSPNs are consistently silent during behaviors, a property that is not observed for dSPNs.

**Fig. 5:**
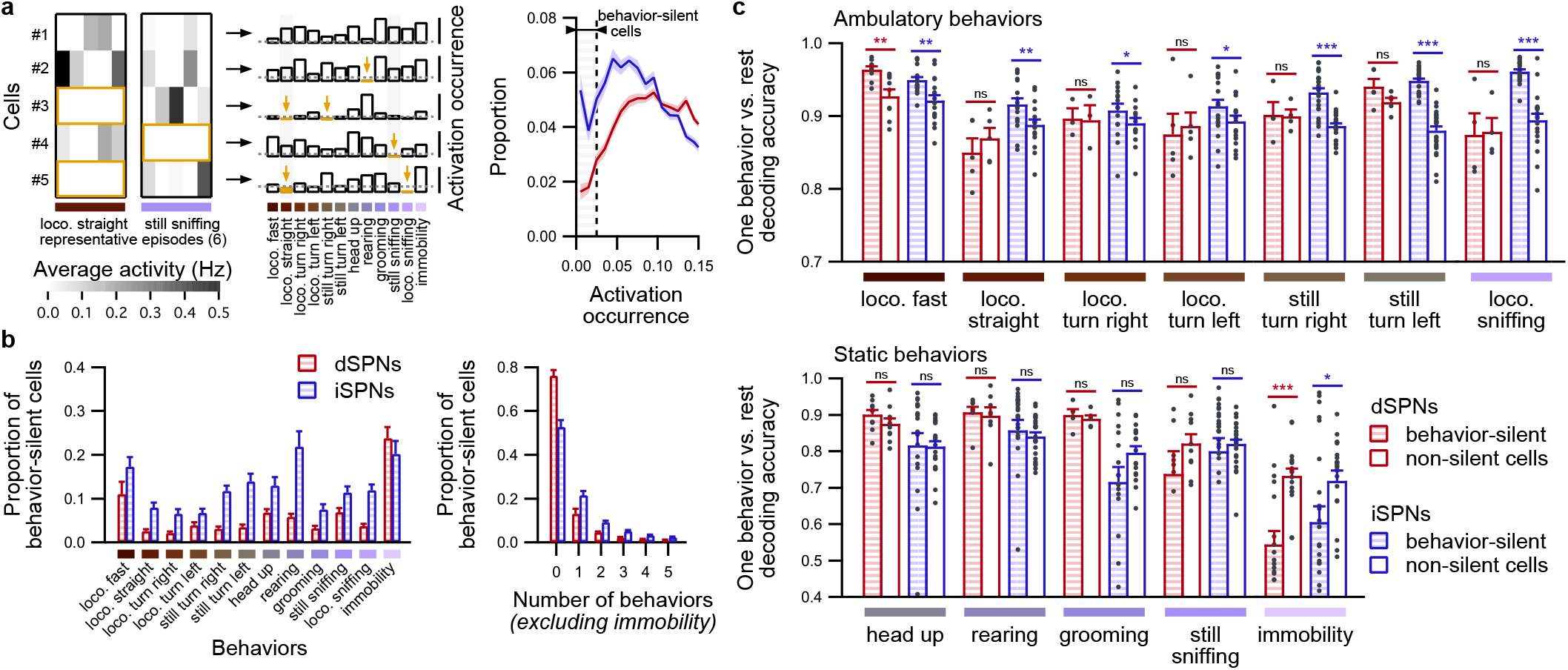
Relevant information for decoding behaviors lies primarily in silent iSPNs. **a**, Behavior-silent cells are identified according to how often these cells are active during episodes of a given behavior, as illustrated by 5 representative neurons (left panel; 6 representative episodes displayed) that displayed various levels of activation occurrence for each behavior (middle panel). Some neurons are rarely or never active during all episodes of a given behavior (middle panel, yellow bars and arrows). The activation occurrence threshold was set to 0.025 according to the distribution of activation occurrence values for all behaviors in dSPNs and iSPNs (right panel; threshold indicated by the black dashed vertical line). **b**, Average fraction of dSPNs and iSPNs identified as behavior-silent for each behavior (left panel) and average fraction of dSPNs and iSPNs categorized as behavior-silent during no behavior or one to five behaviors (right panel). **c**, One behavior vs. rest decoding accuracy (simple matching coefficient) for prediction in separating each behavior from other behaviors using neurons that were classified as behavior-silent during this behavior (hatched bars) or nonsilent (unfilled bars) in dSPNs (red; n = 29 sessions in 8 mice) or iSPNs recordings (blue; n = 37 sessions in 9 mice) (behavior-silent vs. nonsilent neurons: ns, p >0.05, * p < 0.05, ** p < 0.01, *** p < 0.001). Note that the decoding performance is higher for behavior-silent iSPNs than for nonsilent cells during ambulatory behaviors (top panel), whereas this result is not observed during static behaviors (bottom panels).

### Long-term stability of neural code over weeks

To reinforce our previous findings and assess their reliability, we investigated their long-term stability. Thus, for each animal, we performed a longitudinal registration of neurons^29^ across pairs of recording sessions over one month (Extended Data Figs. 1 and 8a). The registration performance was similar for dSPN and iSPN recordings (Extended Data Fig. 8b). On average, 104 ± 5 cells were recovered between any pair of sessions (Extended Data Fig. 8c), corresponding to an overlap of 33.9 ± 0.8% in the subset of cells identified in both sessions, ranging from 36.2 ± 1.5% after approximately 5-7 days to 32.5 ± 1.4% after approximately 30 days. As described above, we first quantified the neuronal activation similarity for each behavior between pairs of sessions. For all behaviors, the neuronal activation similarity was higher for dSPNs and iSPNs than for their respective shuffles, as evaluated by either random permutations of registered cell pairs or by replacing one cell or each pair with its closest neighbor (Extended Data Fig. 8d,e). These controls provide an additional post-hoc validation of the longitudinal registration procedure. Then, comparisons between dSPNs and iSPNs revealed that, with the exception of immobility, the neuronal activation similarity was consistently higher over time for dSPNs than for iSPNs (Fig. 6a, Extended Data Fig. 8d,e). This result appears to indicate that neuronal activation patterns are preserved better for dSPNs than for iSPNs, which may reflect the previously established bias toward activation in the neuronal code of behaviors in dSPNs.

**Fig. 6:**
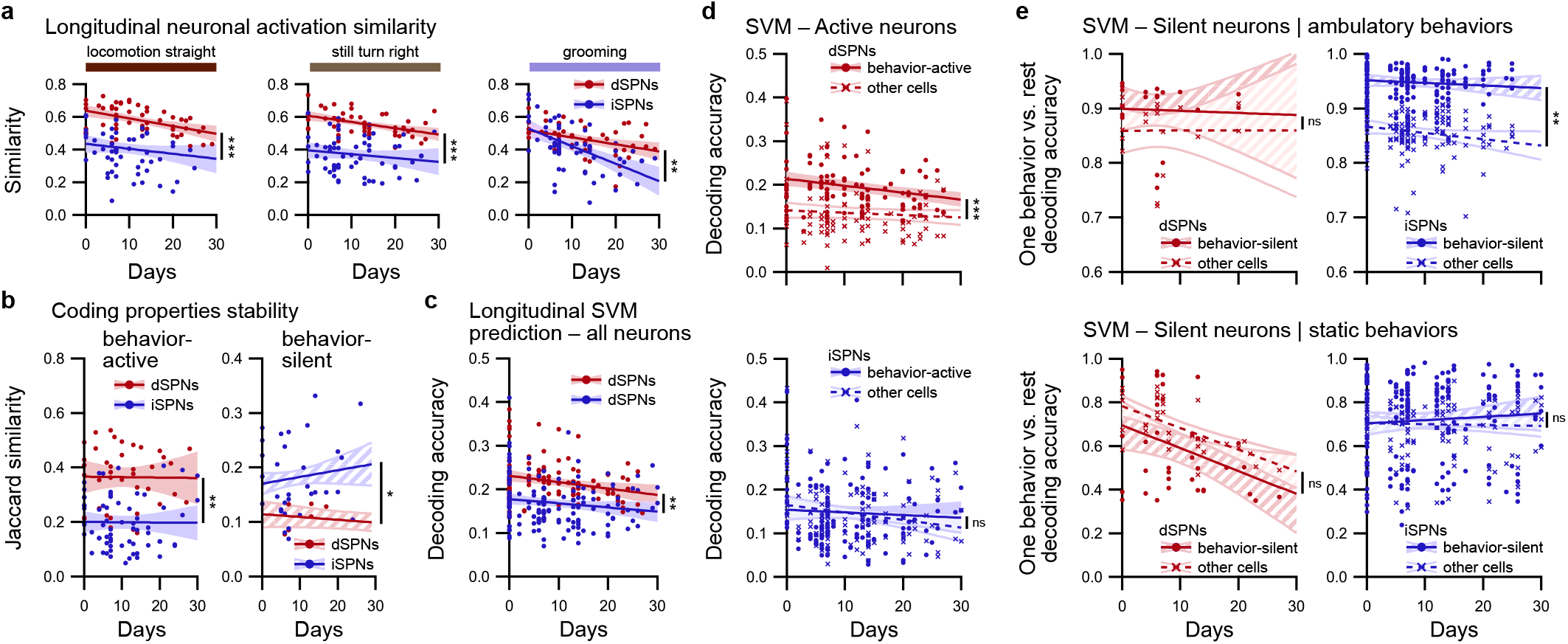
Long-term stability of the respective coding properties of dSPNs and iSPNs. **a**, Differences in neuronal activation similarity during identified behaviors between dSPNs (red; n = 52 pairs of sessions in 8 mice) and iSPNs (blue; n = 62 pairs of sessions in 9 mice) are preserved for weeks, as illustrated for locomotion straight (left), still turn right (middle), and grooming (right) (dSPNs vs. iSPNs: *** p < 0.001). **b**, Quantification of the long-term stability of coding properties of neurons using the Jaccard similarity coefficient between binary classification of longitudinally registered neurons that were labeled as behavior-active (left panel) or behavior-silent during ambulatory behaviors (right panel) in dSPN (red; behavior-active: n = 58 pairs of sessions meeting criterion; behavior-silent: n = 16 pairs of sessions meeting criterion) and iSPN recordings (blue; behavior-active: n = 70 pairs of sessions meeting criterion; behavior-silent: n = 34 pairs of sessions meeting criterion) (dSPNs vs. iSPNs: * p < 0.05, ** p < 0.01). Note that the similarity is higher for behavior-active dSPNs and behavior-silent iSPNs. **c**, Accuracy in predicting the behavior according to the activity of longitudinally registered neurons using SVM classifiers trained on different recording sessions for dSPNs (red; n = 121 pairs of sessions) and iSPNs (blue; n = 171 pairs of sessions) (dSPNs vs. iSPNs: ** p < 0.01). **d**, Decoding accuracy for longitudinal predictions of behaviors using behavior-active neurons (circles, plain line, colored confidence interval) or non-behavior-active cells (crosses, dashed line, unfilled confidence interval) in dSPN (top panel; red; n = 93 pairs of sessions meeting criterion) or iSPN recordings (bottom panel; blue; n = 132 pairs of sessions meeting criterion). The accuracy remained higher for behavior-active dSPNs than non-behavior-active dSPNs for many days, whereas the accuracy was similar for both groups of iSPNs (behavior-active vs. other cells: ns p > 0.05, *** p < 0.001). **e**, Simple matching coefficient for long-term predictions on separating one behavior from other behaviors using neurons classified as behavior-silent (circles, plain line, hatched confidence interval) or nonsilent (crosses, dashed line, unfilled confidence interval) during this behavior and pooled for ambulatory behaviors (top panels) and static behaviors (bottom panels) for dSPNs (left panels; red; ambulatory: n = 18 pairs of sessions meeting criterion; static: n = 46 pairs of sessions meeting criterion) and iSPNs (right panels; blue; ambulatory: n = 52 pairs of sessions meeting criterion; static: n = 69 pairs of sessions meeting criterion). Note that the only difference between behavior-silent and non-behavior-silent cells is observed for iSPNs during ambulatory behaviors (behavior-silent vs. other cells: ns p > 0.05, ** p < 0.01).

Following this idea, we evaluated the stability of the behavior-coding properties across sessions by comparing the classification of cells into the behavior-excited or behavior-silent categories using the Jaccard index (intersection over union) for binary attributes. We observed that within registered cells between pairs of sessions, the proportion of neurons that remained classified as behavior-active for the same behaviors was higher among dSPNs than among iSPNs (Fig. 6b). Conversely, during ambulatory behaviors, behavior-silent iSPNs overlapped between sessions more often than behavior-silent dSPNs (Fig. 6b, Extended Data Fig. 9c). This result indicates that the classification of dSPNs as behavior-active and the classification of iSPNs as behavior-silent are more consistent over time than their respective counterparts in the other striatal population.

To deepen this observation on the categorization of single neurons and extend it to neural ensembles, we quantified whether support vector machines trained on neuronal activity and behavior time series on a given day can efficiently predict the behavior using the neuronal activity of the same neurons on a different day. For both dSPNs and iSPNs, the longitudinal prediction of behaviors using support vector machines is consistently more accurate than the predictions obtained with classifiers trained on time-lagged data (Fig. 6c, Extended Data Fig. 9a,b). Interestingly, the longitudinal prediction of behaviors performs better with dSPNs than with iSPNs, which might reflect the higher long-term stability of neuronal activation similarity for dSPNs.

Finally, we analyzed whether the activity of dSPNs and iSPNs categorized as behavior-active or behavior-silent on a given day can be used to efficiently predict behaviors on a different day. We observed that for dSPNs, behavior predictions were significantly more accurate when only behavior-active cells were used than when non-behavior-active cells were used (Fig. 6d). In contrast, the accuracy is similar when behaviors are predicted with either behavior-active iSPNs or non-behavior-active iSPNs (Fig. 6d). This finding indicates that the bias of the neural code toward behavior-active dSPNs and the information content of these cells in response to behaviors remain conserved for weeks. For behavior-silent neurons (or behavior-inactive neurons, using the alternate definition), we noted that the prediction of ambulatory behaviors is significantly more accurate with behavior-silent iSPNs than with non-behavior-silent cells (Fig. 6e, Extended Data Fig. 9d), whereas no difference was observed for static behaviors. On the other hand, there is no difference in the prediction accuracy when either behavior-silent or non-behavior-silent dSPNs are used (Fig. 6e, Extended Data Fig. 9d). This finding highlights the long-term preservation of the bias toward behavior-silent iSPNs in the striatal neural code. Overall, these results demonstrate the long-term stability of the neuronal encoding properties established in individual recording sessions, reinforcing the reliability of our findings.

Thus, our findings demonstrate that neuronal representations of spontaneous self-paced behaviors in striatal ensembles are biased toward activation in dSPNs and silencing in iSPNs and that these representations are preserved for weeks. These results provide new original insights into both the neuronal organization of striatal ensembles for behavior encoding and efficient motor program selection. In addition, our observations allow us to propose an updated model for behavior encoding in the two striatal pathways that solves discrepancies raised by previously formulated hypotheses.

## DISCUSSION

Neurons in the dorsal striatum exhibit diverse and heterogeneous responses to different external variables^11, 13, 14, 30^. These responses cannot be easily interpreted and highlight the challenge of identifying the distinct encoding properties of the two parallel efferent pathways in the dorsal striatum. In this study, we evaluated the neuronal encoding properties of SPN ensembles during self-paced natural behaviors in an open field. In this experimental paradigm, animals can express unconstrained naturalistic behaviors forming a large behavioral repertoire. In our experiments, we observed previously unreported differences between direct and indirect pathway SPNs. These differences most likely reflect the higher variability of iSPNs in neuron ensembles that are activated during different episodes of the same behavior, which could be observed only during long sessions of open-field exploration. However, despite these heterogeneous response properties, the behaviors could be reliably decoded from the activity of either dSPNs or iSPNs, demonstrating that both pathways contain the same level of information with respect to the ongoing behaviors. Therefore, it questions the respective organization of the neural code in both pathways.

Many formulations have been proposed to explain the organization of neuronal activity in the two striatal pathways. Among them, a “complete selection-suppression” model proposes prokinetic and antikinetic functions for the direct and indirect pathways, respectively^4–6, 17^ (Extended Data Fig. 10a). This model proposes that proper motor program execution relies on the congruent activation of a small discrete subpopulation of dSPNs that encode this motor program and a large number of iSPN ensembles associated with all other motor programs that inhibit all other behaviors. Such a model predicts that dSPNs are more selective for actions than iSPNs, which is inconsistent with observations in previous reports^11, 14–16^ and our results. However, this model is compatible with our observation that the neural code among dSPNs is biased toward activation, whereas the neural code is biased toward inhibition in iSPNs because during a given behavior, dSPNs associated with this behavior are activated, whereas iSPNs associated with the same behavior remain silent (Extended Data Fig. 10d). On the other hand, the “cooperative selection” model^13^ (Extended Data Fig. 10b) proposes the cooperative activation of discrete dSPNs and iSPNs ensembles that are similarly tuned toward behaviors to select proper motor programs. Although this model accurately incorporates the coactivations of similar size dSPN and iSPN ensembles during actions, it does not account for the functional dissimilarities of dSPNs and iSPNs^4–6^ or our observations of dissimilar encoding dynamics between the two striatal pathways, in particular the bias toward silencing in iSPNs (Extended Data Fig. 10d). As a result, the current models of striatal organization must be reevaluated. This reevaluation needs to take into account our novel observations. Here, we report for the first time divergent behavior-encoding properties between dSPNs and iSPNs, which may reflect their functional dissimilarity. In particular, our observation that the same dSPNs are more consistently activated for each behavior, whereas iSPNs activation is more inconsistent, reveals that behavior specificity is milder in iSPNs than in dSPNs. This consideration is compatible with the general framework of the “complete selection-suppression” model. Additionally, one aspect of our recordings that should be considered is the high variability in neuronal activation in both dSPNs and iSPNs for different episodes of the same behavior. This feature has not been integrated into current models and likely reflects the large effect of external and internal contexts for all occurrences of a given motor program. We thus propose a novel formulation we call “adaptive selection-suppression” (Extended Data Fig. 10c), which attempts to reconcile the abovementioned models, previously reported observations, and our observations (Extended Data Fig. 10d). We hypothesize that dSPNs encode a small subset of accessible behaviors in the overall behavioral repertoire, including the observed behavior, and are highly dependent on the ongoing context and activated to promote these behaviors. Furthermore, specific iSPNs that encode competing behaviors, which are also highly dependent on the context, are activated to suppress these competing behaviors. In the subsequent basal ganglia nuclei, the comparison between selecting dSPN activations and suppressing iSPN activations dictates the ongoing motor program. This model supports our observation that similar behaviors correspond to comparable neuronal activations in dSPNs and iSPNs, as adjacent, highly similar behaviors are more likely to compete with each other than considerably different behaviors. Moreover, an important feature of the neural code in this model is that specific subgroups of dSPNs are activated in response to the expressed motor program, whereas specific subgroups of iSPNs are consistently inactive. This finding is substantiated by our observation that the most relevant information for predicting behaviors is located in dSPNs that are activated during behaviors and iSPNs that remain silent during behaviors. This model guarantees the coactivation of discrete subsets of dSPNs and iSPNs during each behavior and supports the broad prokinetic and antikinetic functions of the direct and indirect striatal pathways, respectively.

As described above, for each observed motor program, our model incorporates the notion of interepisode variability, which depends on the context dictated by both internal and external factors. Indeed, the dorsal striatum incorporates different representations of various contextual modalities, such as spatial information^30, 31^, visual and tactile cues^32^, timing^33^, rewarding and aversive drives^34^, task constraints^15^, and sleep drive^24^. These studies substantiate the idea of rich, highly context-dependent behavior representations supported by SPNs. These representations likely originate in cortical areas^35, 36^ and are transferred to the dorsal striatum for subsequent processing and integration^37–39^. Our model proposes that dSPNs are activated not only in response to the expressed behavior but also in response to additional, often similar behaviors. The concurrent activation of specific iSPN ensembles enables the proper filtering of only one action, most likely the most appropriate action in a given context. This mode of organization likely supports efficient shifts in the behavioral strategy depending on an animal’s internal state or task contingencies^40^.

In addition, in our study, we did not observe a bias toward silencing among iSPNs during static behaviors. This could be because these behaviors contain supplemental substates that we could not separate with our analysis pipeline. Alternatively, this could be a consequence of different modes of encoding in iSPNs during ambulatory and static behaviors, similar to spatial mapping in the dorsal hippocampus, which is prominent mainly during locomotion^27, 28^. This alternation between encoding modes may rely on functional interactions between the dorsal hippocampus and the dorsal striatum^41, 42^.

Furthermore, we demonstrated the long-term stability of the neural code in SPNs. We observed that behavior-related information was retained for weeks in behavior-active dSPNs and behavior-silent iSPNs, reinforcing our findings. More precisely, behavior-coding ensembles display a fluctuating membership that ultimately preserves behavior-related information. This finding supports the idea of stable behavioral representations in the dorsal striatum that exhibit some day-to-day fluctuations at the single-cell level. This coding turnover could preserve a certain level of flexibility within the system, which may enable the formation of new traces for encoding similar motor programs that occur in different environments or varied contexts. This mechanism could be critically engaged during procedural or episodic memory formation^43^.

In summary, we identified clear functional differences between SPNs in the direct and indirect pathways in the dorsal striatum in response to self-paced spontaneous behaviors. Our data indicate that ongoing behaviors can be decoded based on dSPN and iSPN activity patterns, despite the fact that these patterns resemble those associated with similar behaviors. Behavior-specific firing and silencing are more prominent in dSPNs and iSPNs, respectively. Our observations are consistent with a model in which dSPN activations represent the ongoing behavior alongside competing motor programs, while iSPNs specific for the ongoing behavior become silent and iSPNs specific for competing behaviors become active. These observations are critical for a deeper understanding of striatal functional organization and strengthen the view that direct and indirect pathways cooperatively orchestrate motor programs in a manner that is highly dependent on the ongoing context by selecting a small subset of accessible behaviors while suppressing competing behaviors.

## ACKNOWLEDGMENTS

We thank Dr. Philippe Faure and Dr. Clément Léna for their critical reading and their helpful insights on our manuscript. C.V. is a postdoctoral researcher of FRS-FNRS. A.C. is a research fellow of FRS-FNRS. P.B. is a research associate of the FRS-FNRS. A.K.E. is a research director of the FRS-FNRS and a WELBIO investigator. This research was supported by grants from FRS-FNRS (grants #23587797, #33659288, #33659296), WELBIO (#30256053), Fondation Simone et Pierre Clerdent (2018 Prize), and Fondation ULB to A.K.E., and grants from FRS-FNRS (grants #34793348, #35285205) to P.B. The funders had no role in study design, data collection and analysis, preparation of the manuscript, or decision to publish.

## AUTHOR CONTRIBUTIONS

C.V., P.B., and A.K.E. designed the study. A.C. and D.H. deployed in vivo calcium imaging protocols and performed experiments. C.V., A.C. and A.K.E. conceptualized the analyses. C.V. and A.C. established and conducted calcium signal extraction and behaviors identification. C.V. performed the other analyses. A.K.E. supervised the project. C.V., A.C., and A.K.E. wrote the manuscript. All authors discussed the manuscript.

## COMPETING INTERESTS

The authors declare no competing interests.

## DATA AVAILABILITY

The data supporting the findings are available within the article and its supplementary materials.

## EXTENDED DATA FIGURES

**Extended Data Fig. 1:**
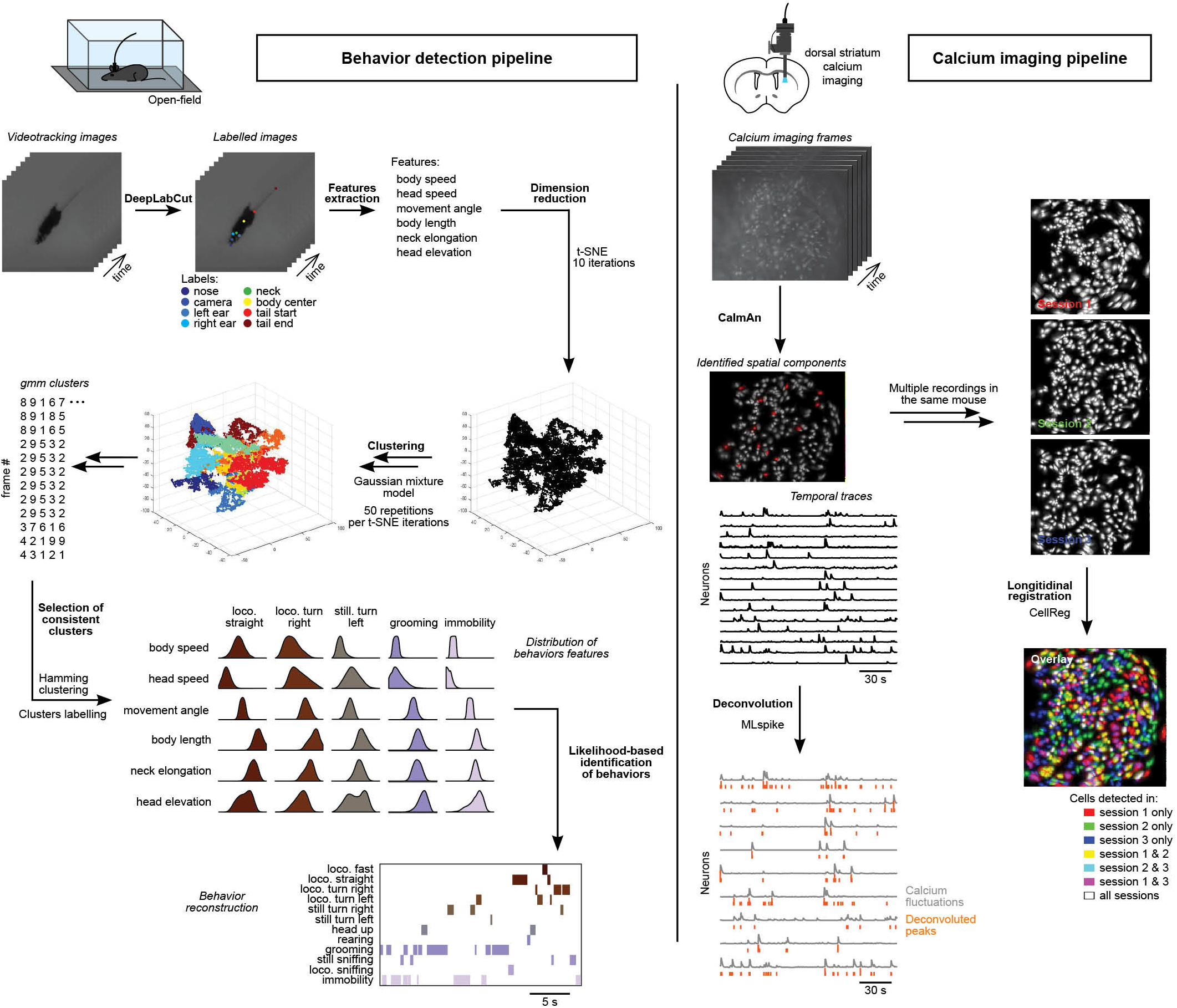
General pipeline for behavioral and calcium data extraction. The synchronous video tracking and microendoscope recordings were analyzed as follows. For video tracking recordings, images are first processed using DeepLabCut to identify and track the position of 8 body parts (nose, camera, ear left, ear right, neck, body center, tail start, and tail end). Using these points, 6 features describing the animal’s posture are computed and fed into multiple iterations of t-SNE dimension reduction and clustering using a Gaussian mixture model. The resulting set of clusters is clustered (Hamming distance) to identify and isolate consistent behavioral clusters and define their corresponding distributions in the feature space. These distributions are used to label all video tracking frames using likelihood-based estimators. In parallel, calcium imaging videos are first processed using CaImAn to separate and identify neurons and extract the temporal evolution of the calcium signal. This signal is then deconvolved using the MLspike algorithm. The cells that are identified during different recording sessions in the same mouse are aligned and longitudinally registered using CellReg.

**Extended Data Fig. 2:**
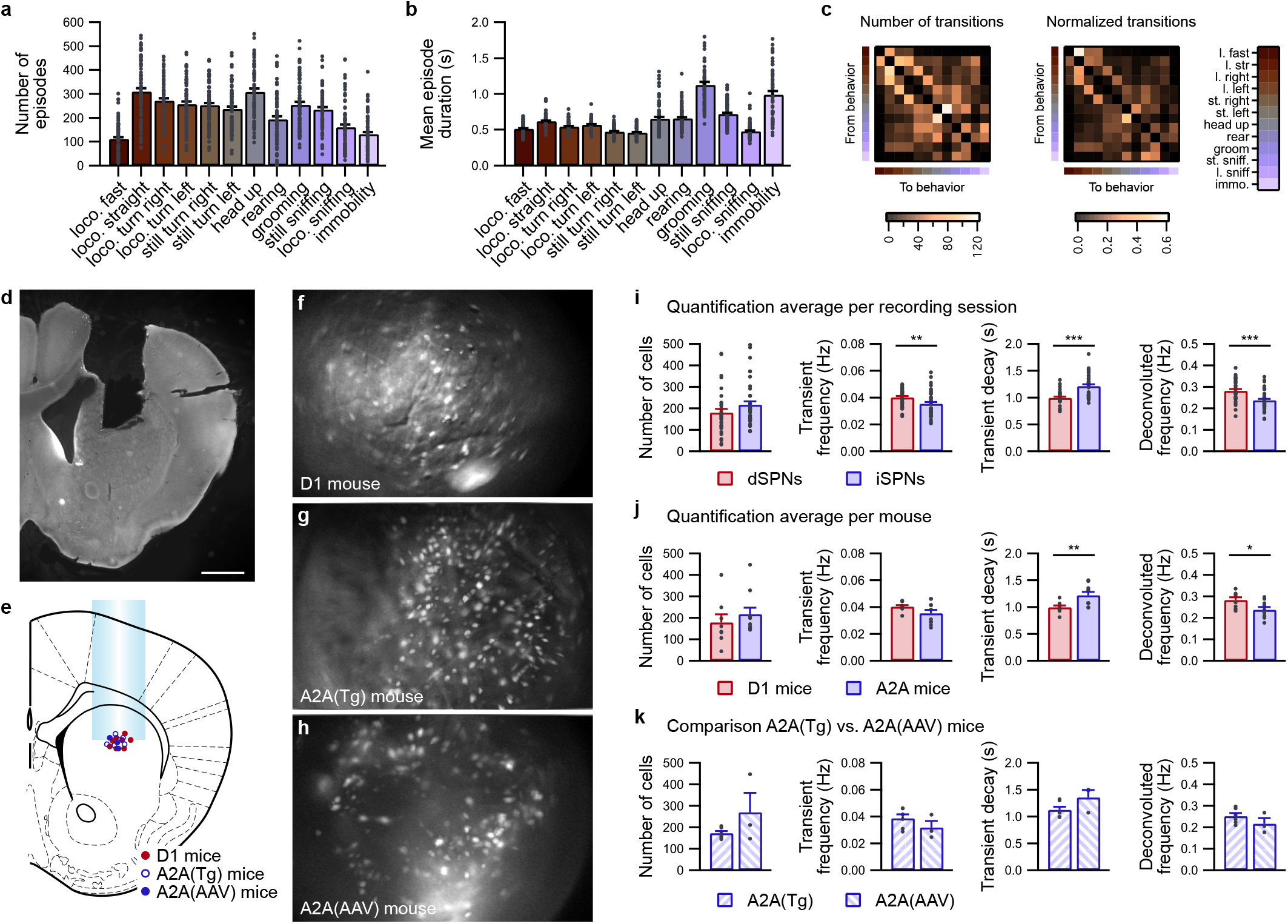
Mice behavior in the open-field and one-photon calcium imaging analyses. **a-c**, Description of mouse behavior architecture during 30 min of undisturbed exploration in an open field (n = 73 sessions in 17 mice), evaluated according to the number of episodes of each behavior (**a**), the average duration of behavior episodes (**b**), and the sequential organization of behaviors, as illustrated by the averaged matrix of the number of transitions (**c**, left panel) from one behavior (lines) to a different behavior (columns), and the same normalized matrix (**c**, right panel), to evaluate the probability when stopping one behavior to begin any of the other 11 behaviors. **d**, Photomicrograph of a coronal section from the brain of a mouse implanted with a GRIN lens in the striatum. **e**, Reconstruction of the GRIN lens positions for all D1 mice (n = 8) and A2A mice (A2A(Tg), n = 6; A2A(AAV), n = 3). **f-h**, Representative image of the maximum fluorescence intensity projection of SPNs labeled with GCaMP6s in a D1 mouse (**f**), A2A(Tg) mouse (**g**), and A2A(AAV) mouse (**h**). **i**, Quantification for each recording session of the identified neuron number, calcium transient average frequency, average transient decay characteristic time, and deconvolved calcium activity for dSPNs (red; n = 33 sessions in 8 mice) and iSPNs (blue; n = 40 sessions in 9 mice) (dSPNs vs. iSPNs: ** p < 0.01; *** p < 0.001). **j**, Same as **i**, averaged per mouse (D1 mice, n = 8; A2A mice, n = 9) (D1 vs. A2A: * p < 0.05; ** p < 0.01). **k**, Comparison of the above parameters, targeting GCaMP6s expression in iSPNs using transgenic reporter mice (A2A(Tg), n = 6) or AAV injection (A2A(AAV), n = 3). No significant difference was observed between A2A(Tg) mice and A2A(AAV) mice.

**Extended Data Fig. 3:**
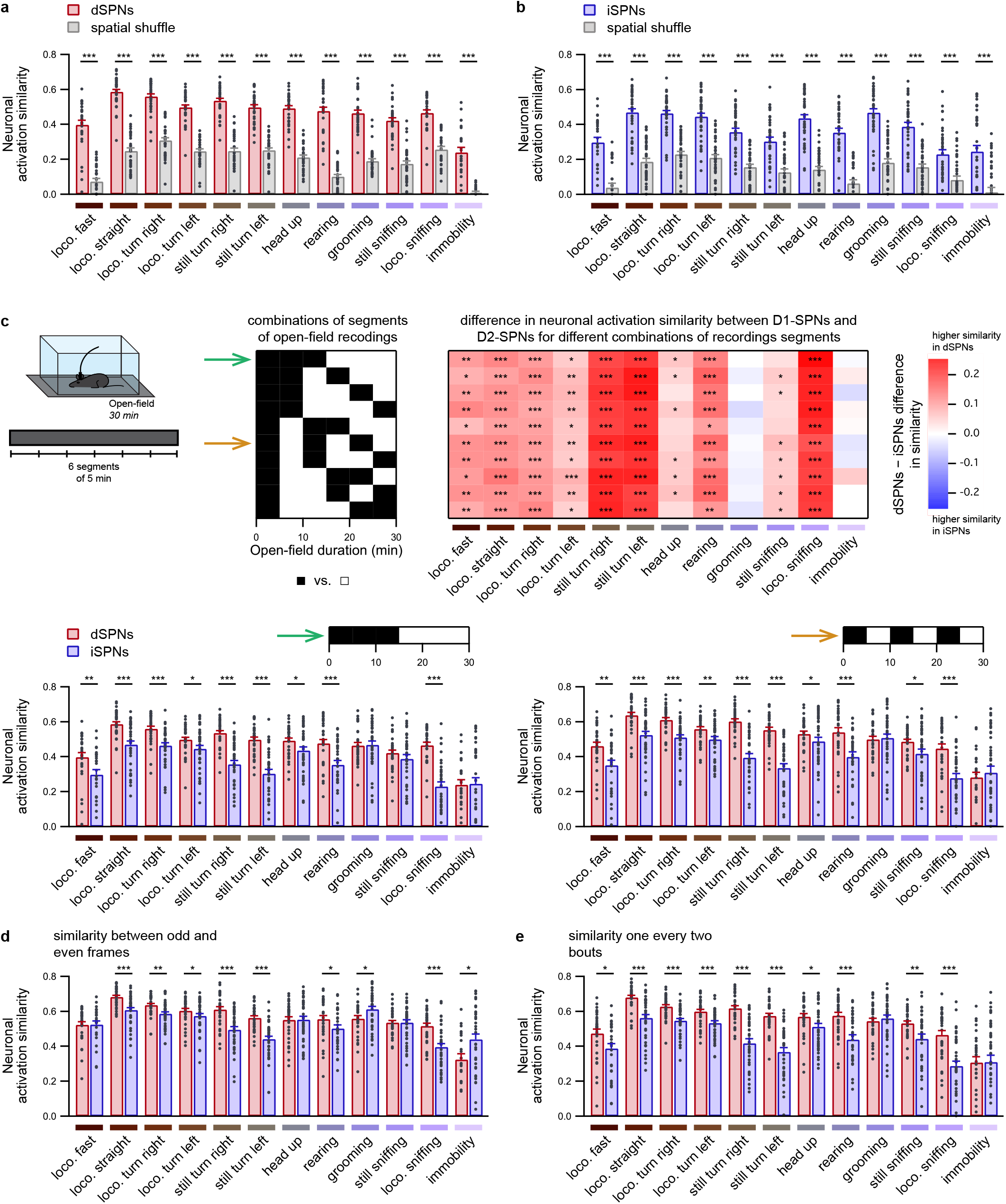
Time invariance of differences in neuronal activation similarity between dSPNs and iSPNs. **a-b**, Neuronal activation similarity between the first and second halves of open-field exploration in dSPNs (**a**) (red; n = 33 sessions in 8 mice) and iSPNs (**b**) (blue; n = 40 sessions in 9 mice) compared with the spatial shuffle of neurons between the first and second halves of open-field exploration (gray bars) (dSPNs or iSPNs vs. shuffle: *** p < 0.001). **c**, The 30 min open-field recording is split into 6 segments lasting 5 min each (top left panel), and the difference between the dSPN and iSPN neuronal activation similarities is calculated for all combinations of 5 min segments into two 15 min segments (3 segments per groups) (top middle and right), highlighting that the difference in similarity between dSPNs and iSPNs is a time-invariant property of the neuronal code. (Bottom) Detailed representation of neuronal activation similarity for two splitting schemes, indicated by colored arrows (dSPNs vs. iSPNs: * p < 0.05, ** p < 0.01, *** p < 0.001). **d**, Neuronal activation similarity between dSPNs (red; n = 33 sessions in 8 mice) and iSPNs (blue; n = 40 sessions in 9 mice) for each behavior, as computed by comparing odd and even frames (dSPNs vs. iSPNs: * p < 0.05, ** p < 0.01, *** p < 0.001). **e**, Neuronal activation similarity between dSPNs (red; n = 33 sessions in 8 mice) and iSPNs (blue; n = 40 sessions in 9 mice), as computed by comparing one episode out of two for each behavior to complementary episodes (dSPNs vs. iSPNs: * p < 0.05, ** p < 0.01, *** p < 0.001).

**Extended Data Fig. 4:**
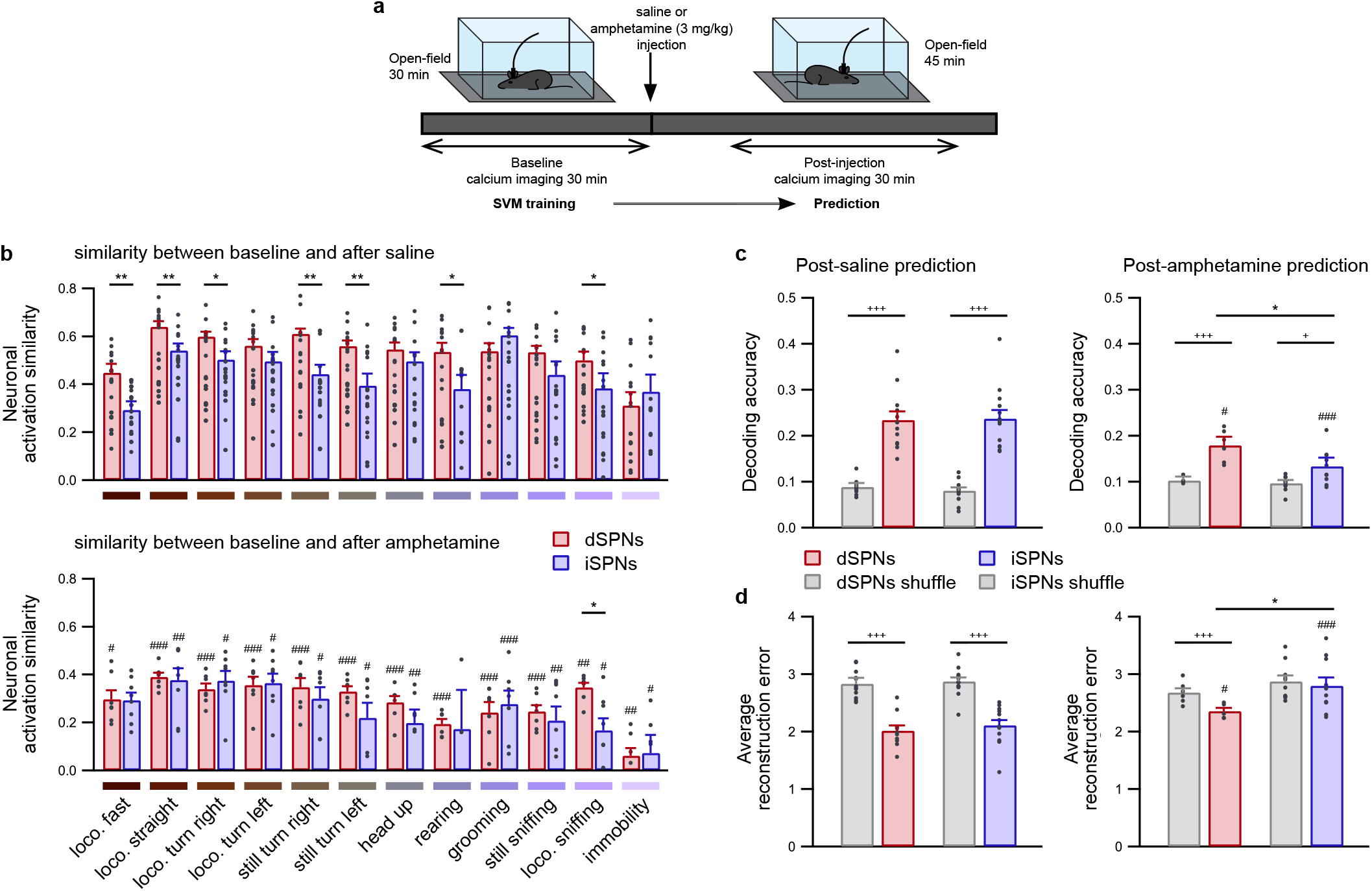
Amphetamine administration reduces the neuronal activation similarity of dSPNs and iSPNs and disrupts SVM-based behavior predictions. **a**, Mice expressing GCaMP6s in either dSPNs or iSPNs freely explored a well-known open field for 30 min (baseline period), received an injection of either saline or amphetamine (3 mg/kg), and were placed back into the open field for an additional 45 min. The post-injection period began 10 min after the injection and lasted 30 min. The SVM classifiers were trained on the baseline period, and the predictions were performed based on the neuronal activity in the post-injection period. **b**, Neuronal activation similarity between baseline and postinjection periods of open-field exploration in dSPNs (red) and iSPNs (blue) for all detected behaviors following saline (top; dSPNs: n = 12 sessions in 8 mice; iSPNs: n = 11 sessions in 9 mice) or amphetamine (3 mg/kg) injection (bottom; dSPNs: n = 7 sessions in 7 mice; iSPNs: n = 8 sessions in 8 mice) (dSPNs vs. iSPNs: * p < 0.05, ** p < 0.01, *** p < 0.001; saline vs. amphetamine: ^#^ p < 0.05, ^##^ p < 0.01, ^###^ p < 0.001). **c-d**, Accuracy of behavior prediction (**c**) and mean behavioral reconstruction error (**d**) using dSPNs (red) or iSPNs (bleu) after saline (left panels; dSPNs: n = 10 sessions in 6 mice; iSPNs: n = 13 sessions in 9 mice) or amphetamine administration (right panels; dSPNs: n = 6 sessions in 6 mice; iSPNs: n = 9 sessions in 9 mice), as compared with time-lagged data (gray). (dSPNs vs. iSPNs: * p < 0.05; saline vs. amphetamine: ^#^ p < 0.05, ^##^ p < 0.01, ^###^ p < 0.001; dSPNs or iSPNs vs. shuffle: ^+^ p < 0.05; ^+++^ p < 0.001).

**Extended Data Fig. 5:**
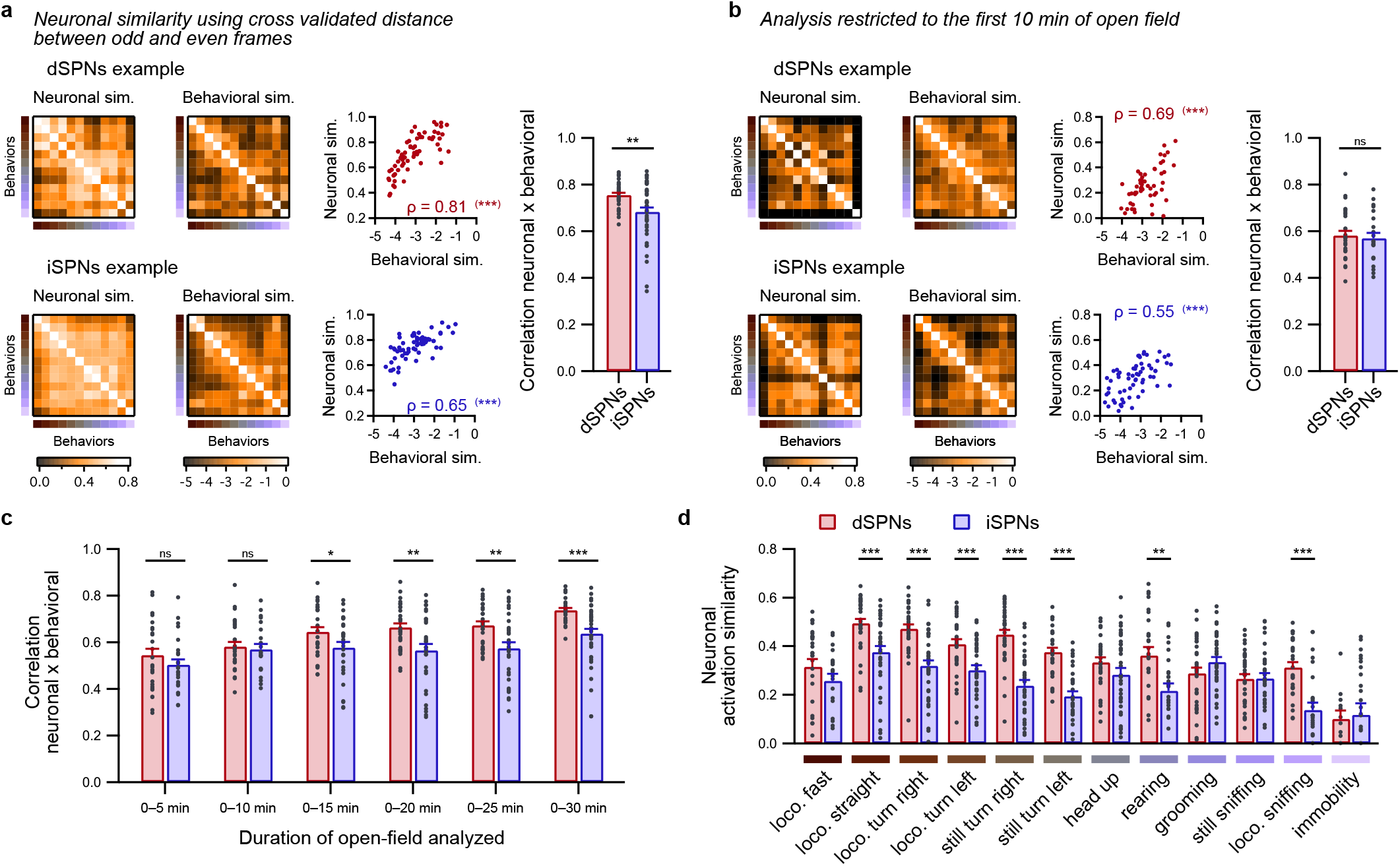
Complements related to correlations between pairwise neuronal similarity and behavioral similarity between behaviors. **a**, Examples of matrices of pairwise neuronal activation similarity between behaviors computed using the cross-validated Euclidean distance between odd and even frames, matrices of pairwise behavioral similarity between behaviors (left panels), and the corresponding scatterplot for pairs of behaviors (middle; Spearman correlation, *** p < 0.001) for one session from one representative D1 mouse (top) and one representative D2 mouse (bottom). The average correlation coefficient (right) between the pairwise behavioral and neuronal similarities was higher for dSPNs (red; n = 33 sessions in 8 mice) than for iSPNs (blue; n = 40 sessions in 9 mice) (dSPNs vs. iSPNs: *** p < 0.001). The results are similar to those displayed in Fig. 2c-f. **b**, Same as **a,** with both the neuronal and behavioral similarities computed for only the first 10 min of open-field exploration. In this condition, the average correlation between the behavioral and neuronal similarities was similar for dSPNs (red; n = 24 sessions in 8 mice) and iSPNs (blue; n = 21 sessions in 9 mice) (dSPNs vs. iSPNs: ns p > 0.05). **c**, Comparison of correlation coefficients between behavioral and neuronal similarities evaluated for different durations of open-field exploration in increments of 5 min for dSPNs (red; n = 33 sessions in 8 mice) and iSPNs (blue; n = 40 sessions in 9 mice) (dSPNs vs. iSPNs: ns p > 0.05; * p < 0.05, ** p < 0.01, *** p < 0.001). **d**, Neuronal activation similarity between the first 5 min and subsequent 5 min of open-field exploration remained higher in dSPNs (red; n = 24 sessions in 8 mice) than in iSPNs (red; n = 21 sessions in 9 mice) during locomotion (loco.) straight, locomotion turn right and left, still turn right and left, rearing, and locomotion sniffing (dSPNs vs. iSPNs: * p < 0.05, ** p < 0.01, *** p < 0.001).

**Extended Data Fig. 6:**
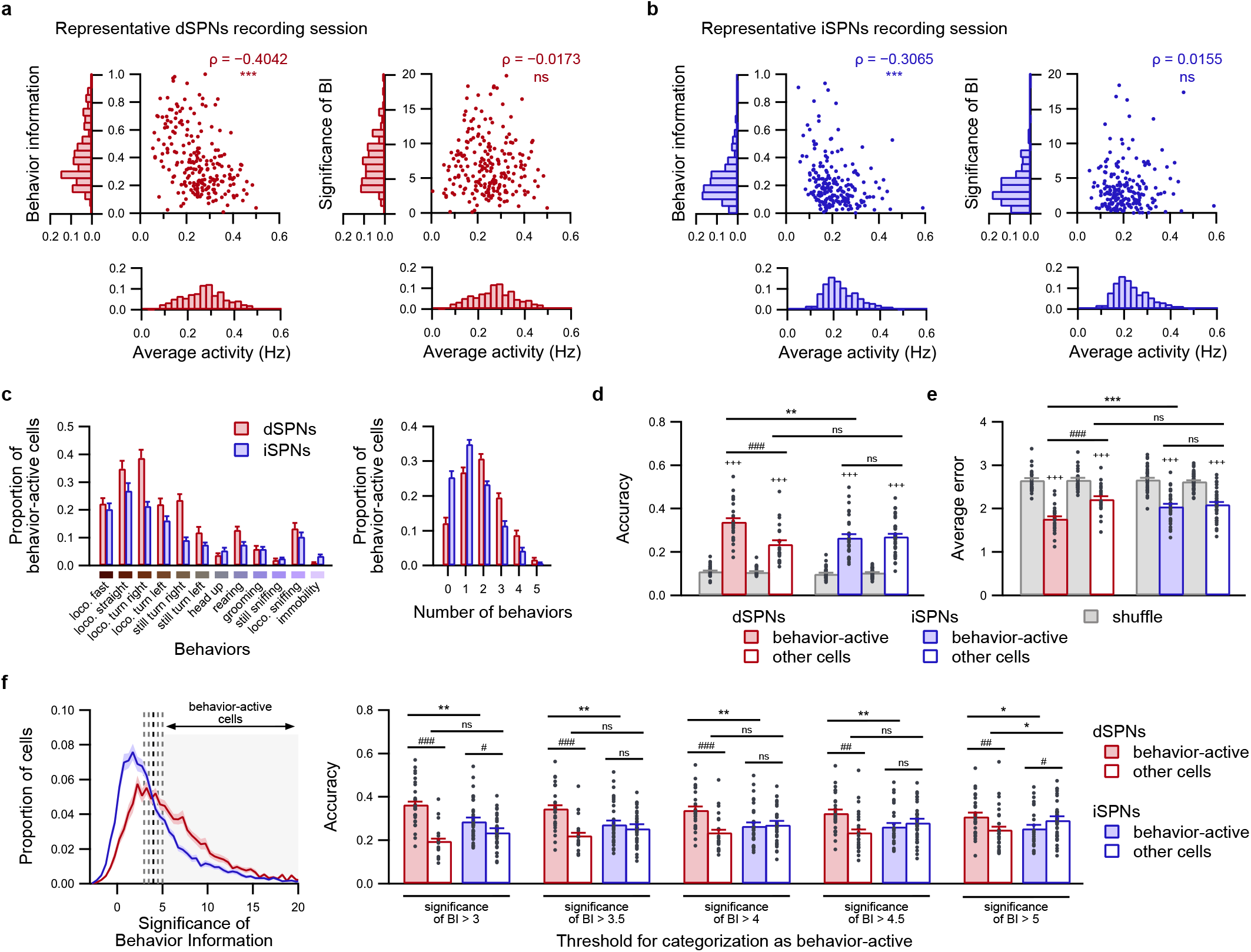
Significance of mutual information and characterization of behavior-active cells. **a-b**, Scatterplots of the relationship between the behavior information (left) or significance of the behavior information (right) and the average activity of dSPNs (**a**) and iSPNs (**b**) in one representative session. Note the significant inverse correlation (Spearman correlation: ns p > 0.05, *** p < 0.001) between the behavior information and event rate, which indicates that the behavior information is biased toward neurons firing at a low rate. Using the significance of the behavior information, which is computed by comparing the observed value of the behavior information to random shuffles, removes this bias. **c**, Average fraction of dSPNs and iSPNs identified as behavior-active (significance of behavior information above 4) for each behavior (left panel) and the average fraction of dSPNs and iSPNs labeled as behavior-active during no behavior or one to five behaviors (right panel) (dSPNs: n = 29 sessions from 8 mice; iSPNs: n = 37 sessions from 9 mice). **d-e**, Decoding accuracy (**d**) and average reconstruction error (**e**) of SVM classifiers applied to behavior-active (plain bars) or non-behavior-active (unfilled bars) dSPNs (red; n = 29 sessions in 8 mice) or iSPNs (blue; n = 37 sessions in 9 mice), compared with the chance level when decoding classifiers trained on time-lagged data (gray bars) (dSPNs vs. iSPNs: ns p > 0.05, *** p < 0.01; behavior-active vs. other cells: ^#^ p < 0.05, ^###^ p < 0.001 observed data vs. time-lagged: ^+++^ p < 0.001). Note that the decoding performance based on either behavior-active neurons or other cells is better than that based on time-shuffled data, and the decoding performance is better with behavior-active cells than with non-behavior-active cells in dSPN recordings (red bars; n = 29 sessions from 8 mice), whereas performance is similar between both cell groups in iSPNs recordings (blue bars; n = 37 sessions from 9 mice). Moreover, the decoding performance is better for behavior-active dSPNs than for behavior-active iSPNs. **f**, Effect of modifying the threshold used for the level of significance of behavior information for labelling neurons as behavior-active (left panel) on the decoding accuracy (right panel) when predicting behaviors using behavior-active neurons (plain bars) or non-behavior-active neurons (unfilled bars) in dSPN (red bars; n = 29 sessions from 8 mice) and iSPN (blue bars; n = 37 sessions from 9 mice) recordings (dSPNs vs. iSPNs: ns, p > 0.05, ** p < 0.01; behavior-active vs. other cells: ^#^ p < 0.05, ^###^ p < 0.001). The use of 5 different thresholds (ranging from 3 to 5 SD, corresponding to the position of the actual behavior information in comparison to the distribution of the behavior information calculated from random permutations) maintains significantly higher decoding performance when using behavior-active dSPNs than when using behavior-active iSPNs.

**Extended Data Fig. 7:**
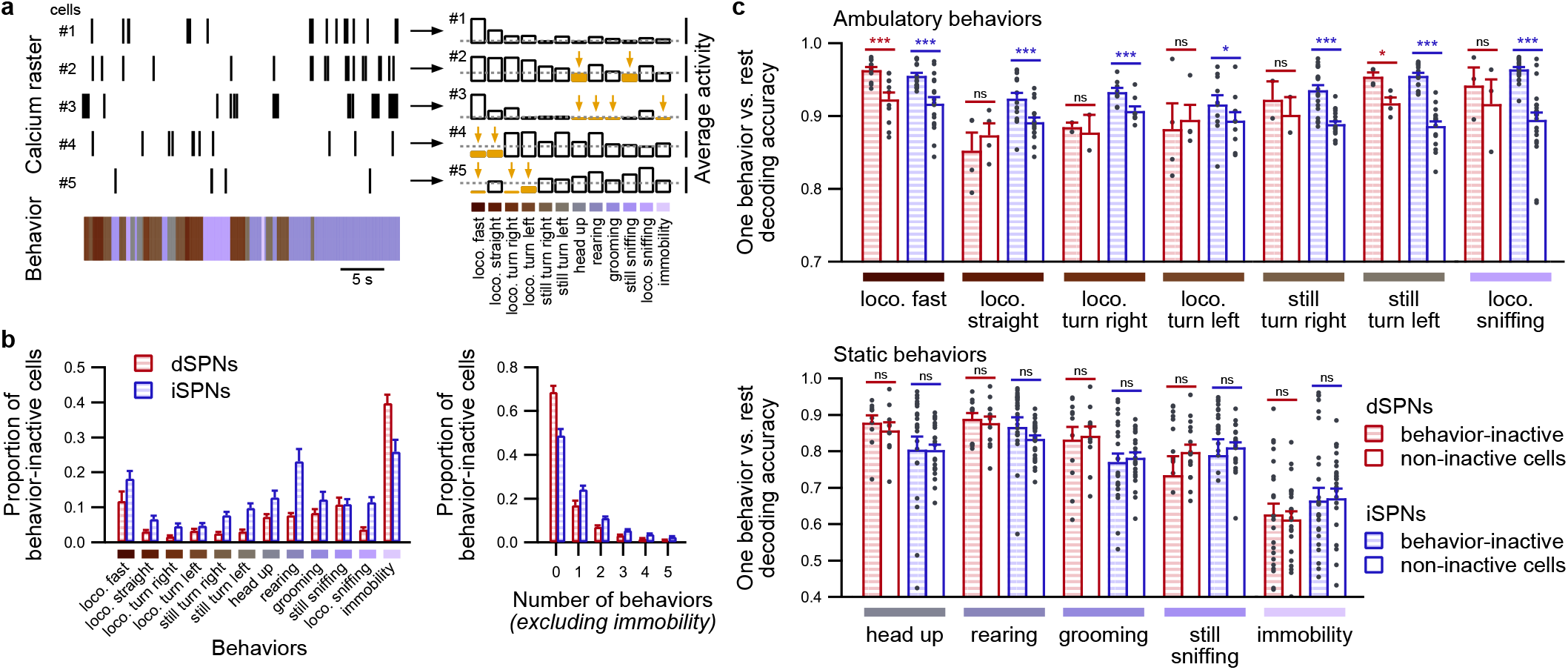
Contribution of behavior-inactive cells to neural code using an alternate definition. **a**, Behavior-inactive cells are identified according to their average activity during each behavior, as illustrated by 5 representative neurons (left panel) displaying different average activities for each behavior (right panel). The threshold for identifying an inhibited cell during a given behavior is set when the cell displays an average activity of less than 0.1 events/s (right panel, yellow bars). **b**, Average fraction of dSPNs and iSPNs labeled behavior-inactive for each behavior (left panel) and average fraction of dSPNs and iSPNs labeled behavior-inactive during no behavior or one to five behaviors (right panel). **c**, Simple matching coefficient for separating each behavior from other behaviors using neurons that are classified as either behavior-inactive during proper behavior (hatched bars) or non-behavior-inactive (unfilled bars) in dSPN (red; 29 sessions in 8 mice) or iSPN recordings (blue; n = 37 sessions from 9 mice) (behavior-inactive vs. non-behavior-inactive neurons: ns, p >0.05, * p < 0.05, ** p < 0.01, *** p < 0.001). The results are similar to those obtained when the neurons are classified as behavior-silent and non-behavior-silent.

**Extended Data Fig. 8:**
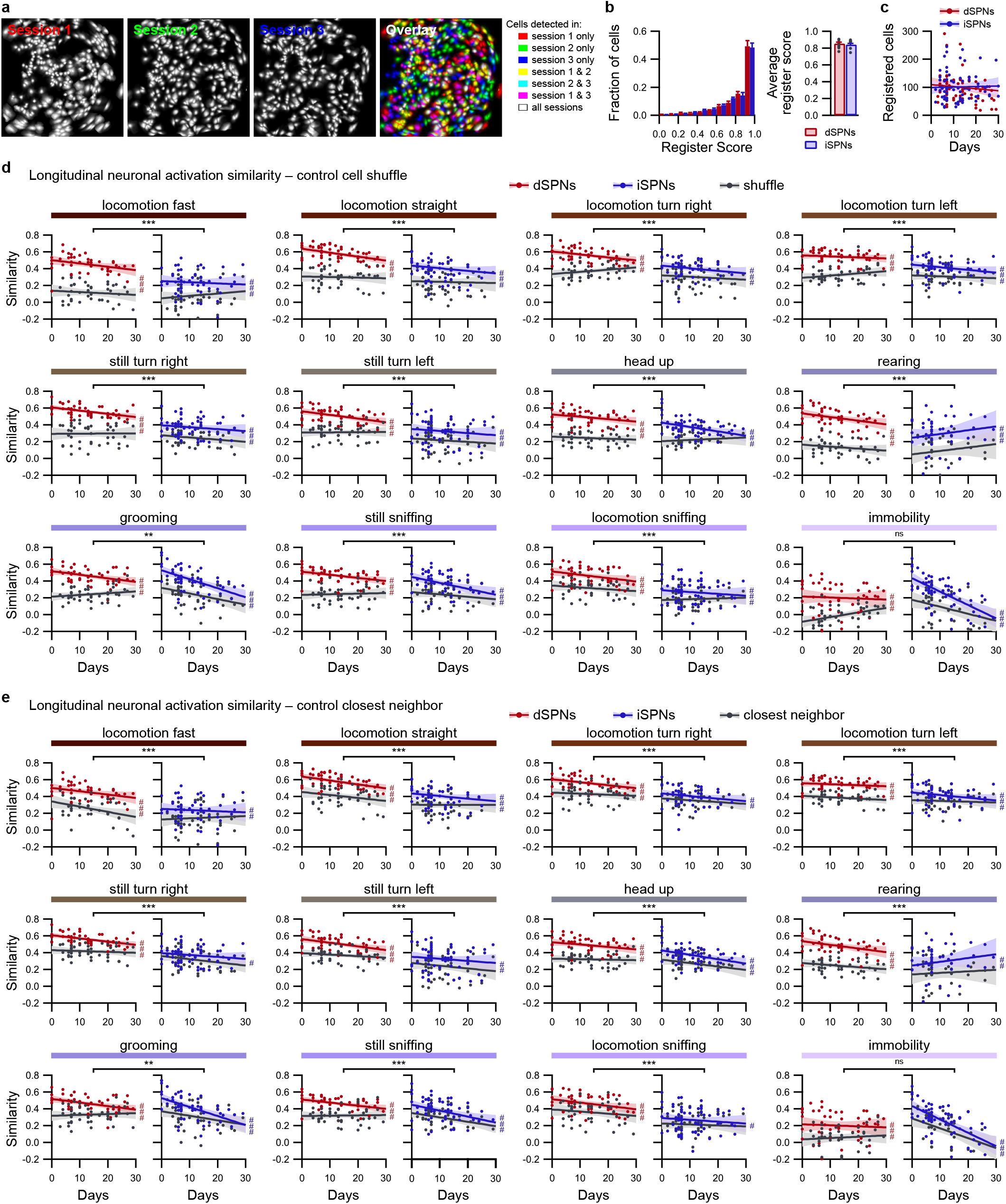
Validation of longitudinal registration of cells and longitudinal evaluation of neuronal activation similarity during behaviors. **a**, Representative example of spatial footprints of neurons detected during three different sessions recorded from the same mouse and an overlay of the aligned spatial footprint maps, which are color-coded according to the sessions during which the cells were detected. **b**, Distribution of register scores between sessions (left panel) and mean register score (right panel) averaged for all D1 (red; n = 8) and D2 (blue; n = 9) mice. **c**, Number of registered cells between pairs of sessions from the same mouse as a function of the number of days between sessions for dSPN (red; n = 72 pairs of sessions in 8 mice) and iSPN recordings (blue; n = 80 sessions in 9 mice). **d-e**, Evolution of the neuronal activation similarity during identified behaviors across days for dSPNs (red; n = 52 pairs of sessions in 8 mice) and iSPNs (blue; n = 62 pairs of sessions in 9 mice) compared to their respective controls (gray), which were obtained by shuffling pairs of registered neurons (**d**) or by replacing one neuron in each pair of registered cells by its spatially closest neighbor (**e**) (dSPNs vs. iSPNs: ns p > 0.05, ** p < 0.01, *** p < 0.001; observed data vs. control: ^#^ p < 0.05, ^##^ p < 0.01, ^###^ p < 0.001).

**Extended Data Fig. 9:**
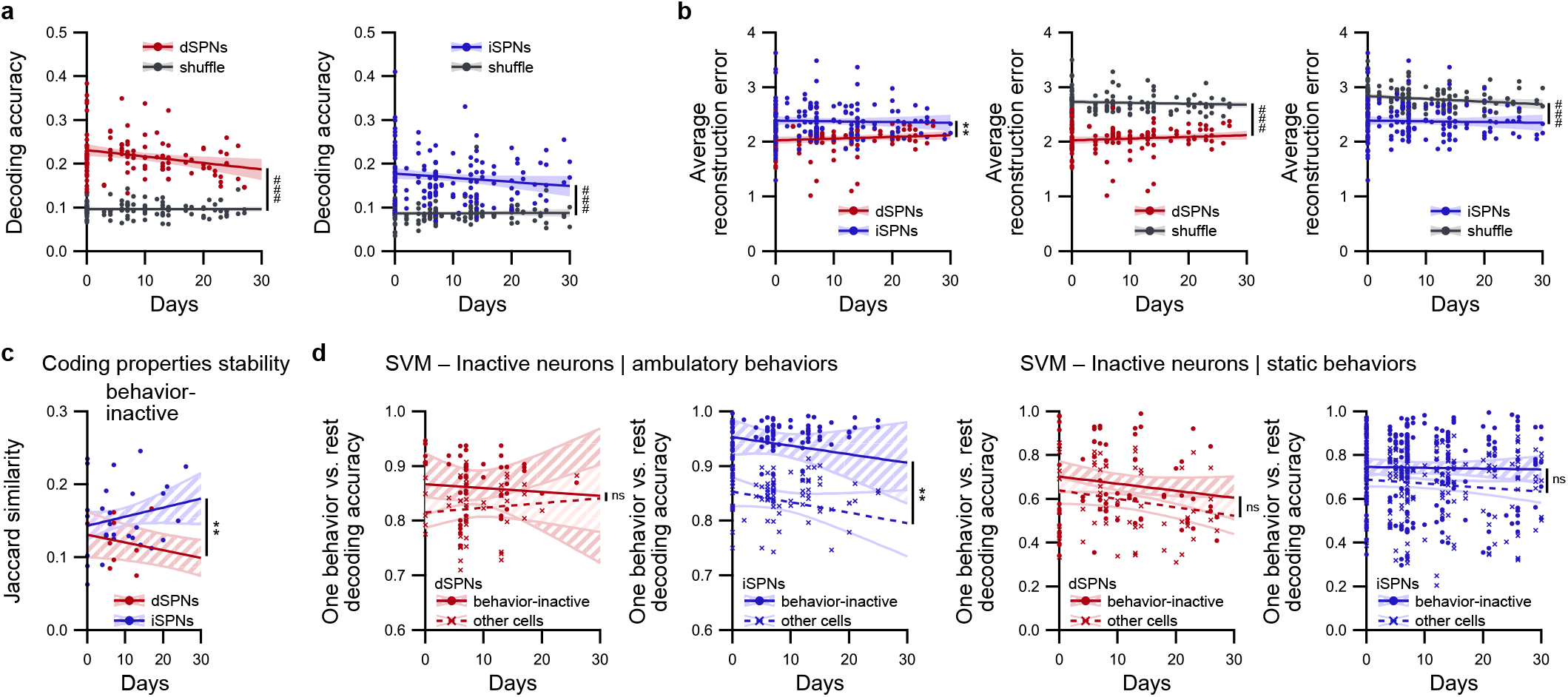
Complements related to longitudinal prediction of behaviors. **a**, Accuracy in predicting the behavior according to the activity of longitudinally registered neurons using SVM classifiers trained on a different recordings for dSPNs (red; n = 121 pairs of sessions) and iSPNs (blue; n = 171 pairs of sessions) and their respective controls, which were obtained using classifiers trained on time-lagged data (gray) (observed data vs. time-lagged: ^###^ p < 0.001). **b**, Average behavioral reconstruction error in predicting the behavior according to the activity of longitudinally registered neurons using SVM classifiers trained on different recordings for dSPNs (red; n = 121 pairs of sessions) and iSPNs (blue; n = 171 pairs of sessions) (dSPNs vs. iSPNs: ** p < 0.01; observed data vs. time-lagged: ^###^ p < 0.001). **c**, Quantification of the long-term stability of the coding properties of neurons using the Jaccard similarity coefficient between the binary classification of longitudinally registered labeled behavior-inactive in dSPN (red; n = 16 pairs of sessions meeting criterion) and iSPN recordings (blue; n = 29 pairs of sessions meeting criterion) (dSPNs vs. iSPNs: ** p < 0.01). **d**, Simple matching coefficient for the long-term prediction of separating one behavior from other behaviors using neurons that were classified as behavior-inactive (circles, plain line, colored confidence interval) or non-behavior-inactive (crosses, dashed line, unfilled confidence interval) during this behavior, pooled for ambulatory behaviors (top panels) or static behaviors (bottom panels) for dSPNs (red; ambulatory: n = 20 pairs of sessions meeting criterion; static: n = 67 pairs of sessions meeting criterion) and iSPNs (blue; ambulatory: n = 46 pairs of sessions meeting criterion; static: n = 75 pairs of sessions meeting criterion) (behavior-inactive vs. non-behavior-inactive: ns p > 0.05, ** p < 0.01). Note that the results obtained for behavior-inactive neurons are highly similar to those observed for behavior-silent neurons.

**Extended Data Fig. 10:**
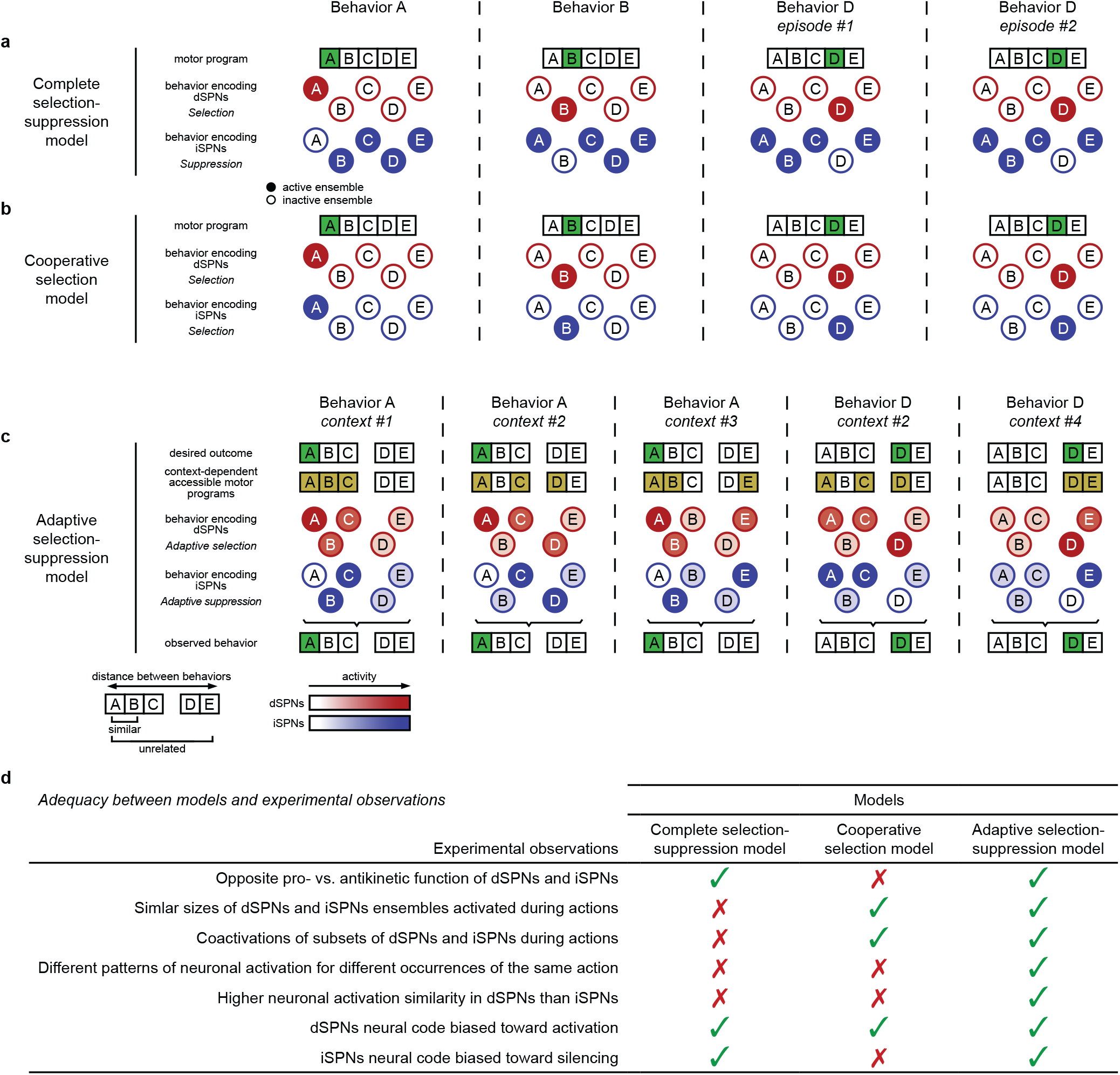
Comparison of theoretical models describing SPN encoding of behaviors. Multiple models have been formulated to describe the organization of the basal ganglia and capture the properties of striatal neurons. **a**, In the “complete selection-suppression” model^17^, subsets of dSPNs and iSPNs are concurrently activated during behaviors. While a specific subpopulation of dSPNs is activated to promote a given motor program, all iSPN subpopulations associated with other motor programs are active to simultaneously suppress these competing motor programs. This model predicts the widespread activation of iSPNs and the focused activation of specific dSPNs. **b**, Alternatively, some models hypothesized a cooperative selection function for dSPNs and iSPNs based on concurrent coordinated activation in both pathways to select proper behaviors^13^. In this model, comparable populations of dSPNs and iSPNs are tuned toward specific behaviors and activated simultaneously to select proper motor programs. This model predicts a high and equivalent level of neuronal activation similarity in dSPNs and iSPNs during episodes of the same behavior, as well as comparable neuronal activation similarities between behaviors in both SPN pathways. **c**, In this study, we propose a new model, referred to here as the “adaptive selection suppression” model. We postulate that at any time, only a few reachable behaviors in the overall behavioral repertoire actual compete with the ongoing/most-desired behavior. These competing behaviors are highly dependent on the ongoing external context (and potentially dependent on the internal state of the animal) and thus differ between different episodes of a given behavior. We hypothesize that, as a result, dSPNs that encode the ongoing behavior and competing behaviors are activated, while dSPNs that encode other behaviors remain silent. At the same time, in the indirect pathway, iSPNs specifically associated with competing behaviors are activated to suppress these competing behaviors, while iSPNs associated with the ongoing behavior remain silent. As a consequence, the activations in the direct and indirect pathways, which select and suppress motor programs, respectively, result in the proper selection of only one ongoing motor program. **d**, When assessing the adequacy between model predictions and experimental observations, we trust that the adaptive selection-suppression model captures more SPN properties, as established in previous studies. First, despite its accurate account of the antagonistic function of dSPNs and iSPNs^4–10^, which implies a bias in the neural code toward activation and inhibition, respectively, the complete selection-suppression model proposes the widespread activation of iSPNs and focused activation of dSPNs during actions, which is not supported by recordings of neuronal activity^11–16^. Additionally, this model cannot explain the differences in neuronal activation similarity we observed in the present study, as this model predicts that populations of neurons that are activated during episodes of the same behaviors are highly similar for dSPNs and iSPNs and almost identical when comparing dSPNs and iSPNs. Furthermore, the cooperative selection model properly captures the simultaneous and targeted activation of subsets of dSPNs and iSPNs during actions^12–14^. However, this model does not explain the antagonistic functions of dSPNs and iSPNs, and because it proposes similar neuronal activation patterns for both dSPNs and iSPNs, it does not account for the differences in the tuning properties in the two pathways we observed in this study. Our proposed novel adaptive selection-suppression formulation aims to reconcile some of the above discrepancies. This model accounts for the prokinetic and antikinetic effects associated with activating dSPNs and iSPNs, respectively. This finding is supported by our observation that the neural code for behaviors is biased toward activation in dSPNs and silencing in iSPNs. Importantly, by incorporating the idea that neuronal activation patterns during self-paced spontaneous exploration of the behavioral repertoire are highly dependent on the current external and internal contexts, we propose that dSPNs and iSPNs activated during different occurrences of the same behavior differ. This result supports the context-dependent variability in activation patterns that we observed using the neuronal activation similarity measure. Moreover, the neuronal clusters in dSPNs that are associated with the ongoing behavior are consistently active, whereas there may be instances when, for the same observed behavior, the sets of activated iSPNs drastically differ (to inhibit different competing behaviors; see, for example, behavior A context #1 vs. behavior A context #3); thus, the proposed model predicts a higher neuronal activation similarity in dSPNs than in iSPNs. In addition, our model incorporates an important feature of the neural code, namely, that in response to the expressed motor program, specific subgroups of dSPNs are activated, whereas specific subgroups of iSPNs are consistently inactive.

## METHODS

### Animal care and use

All procedures were performed according to the Institutional Animal Care Committee guidelines and were approved by the Local Ethical Committee (Comité d’Ethique et de Bien-Être Animal du pôle santé de l’Université Libre de Bruxelles (ULB), Ref. No. 646 N). Mice were maintained on a 12-hour dark/light cycle (lights on at 8 pm) with *ad libitum* access to food and water. The room temperature was set to 22 ± 2 °C with constant humidity (40–60%). The behavioral tests were performed during the dark photoperiod. Both male and female transgenic mice (≥ 8 weeks) were used in all behavioral experiments.

### Transgenic mouse generation

The genetic background of all transgenic mice used in this paper is C57Bl/6J. The mice were heterozygous and maintained by breeding with C57Bl/6 mice. All lines were backcrossed with C57Bl6 mice for at least 8 generations. Three transgenic mouse lines were used: A_2A_-Cre^7^, D_1_-Cre (EY262; GENSAT)^44^ and Ai162/TIT2L-GC6s-ICL-tTA2 (Ai162D; Allen Institute)^45^. Simple transgenic A_2A_-Cre or D_1_-Cre mice (A2A(AAV) and D1 mice, respectively) were used for the virally mediated targeting of iSPNs or dSPNs, respectively. Double transgenic A_2A_-Cre x Ai162/TIT2L-GC6s-ICL-tTA2 mice (A2A(Tg) mice) were generated by breeding and used for targeting iSPNs without virus injection. For this breeding, mice were maintained with doxycycline food pellets (A03 1 g/kg doxycycline hyclate pellet; SAFE, France) to prevent GCaMP6s expression during development and early life stages in offspring. Standard food was introduced when offspring were weaned (3 weeks postnatal). The results of A_2A_-Cre mice expressing GCaMP6s through a virus-mediated strategy (A2A(AAV) mice) or through a double transgenic strategy (A2A(Tg) mice) were compared, and no differences were observed (Extended Data Fig. 2g-h, k). As a consequence, these animals were pooled.

### Viral injections and chronic lens implantation

Under isoflurane anesthesia (induction 4%, maintenance 1% in O_2_; 0.5 L/min), male and female A_2A_-Cre and D_1_-Cre mice (≥ 8 weeks old), which targeted iSPNs and dSPNs, respectively, received two injections (500 nL per site at 100 nL/min) under stereotaxic control in the dorsal striatum (AP: +1.2 mm; ML: –1.75 mm; DV: –3 mm; and AP: +1.2 mm; ML: –1.75 mm; DV: –3.3 mm relative to Bregma) of a Cre-dependent virus encoding GCaMP6s (AAV-Dj-EF1α-DIO-GCaMP6s; Stanford Vector Core, titer 5.02 x 10^12^ vg/ml; UNC Vector Core, titer 3.9 x 10^12^ vg/ml), which was delivered through a cannula connected to a Hamilton syringe (10 µL) placed in a syringe pump (KDS-310-PLUS, KDScientific). Cannulas were lowered into the brain and left in place for 10 min after infusion. Mice (4–5 weeks after virus injection or ≥ 8 weeks old for A2A(Tg) mice) were then prepared for in vivo calcium imaging. A gradient index (GRIN) lens (1.0 mm or 0.6 mm diameter, ID-1050-004605 or ID-1050-004608; Inscopix, Palo Alto, CA) was implanted into the dorsal striatum under stereotaxic control directly above the injection site (AP: +1.2 mm; ML: –1.9 mm; DV: –2.8 mm relative to Bregma). Once the lens was positioned, the lens was secured to the skull using Metabond (C&B, Sun Medical Co. Ltd, Japan) and protected using tape. Two weeks after lens implantation, a microendoscope baseplate (ID-1050-004638; Inscopix) was attached to the skull with Metabond in the optimal imaging plane (550 µm above the lens for the 0.6 mm diameter lens and ∼300 µm above the lens for the 1.0 mm diameter lens). The behavioral experiments began at least one week after the baseplate was fixed to ensure that the field of view was adequately cleared.

### Open-field behavior

The behavior experiments were conducted in an open field arena (40 cm x 40 cm x 40 cm, length x width x height) with white walls in a dark environment (0 lux). Before the first recording session, mice were habituated to the open field and the microendoscope for at least 5 consecutive days by using a dummy microscope (ID-1050-003762; Inscopix) mounted in place of the actual microscope. The animal behavior was recorded for 30 min for 4–5 sessions spaced over 5–7 days using a camera (sampling rate: 40 fps) mounted on the ceiling approximately 1.5 m above the arena controlled through EthoVision XT14 (Noldus). One photon calcium imaging was sampled at 20 fps using a nVista 3.0 microendoscope (Inscopix). A commutator (Inscopix) attached to the ceiling was placed between the camera and the acquisition box to minimize cable entanglement. To prevent photobleaching, calcium frames were acquired with a 2 min OFF / 3 min ON pattern, and time synchronization between the calcium recordings and open-field videos was programmed and managed through EthoVision XT14.

At the end of some recording sessions, mice received an intraperitoneal (i.p.) injection of either saline or amphetamine (3 mg/kg in saline) and were immediately placed back into the open-field arena for an additional 45 min of behavior and calcium imaging following the same acquisition protocol. Amphetamine treatment always occurred on the last day of recording to prevent any effect on neuronal activity due to the long-lasting effects of amphetamine.

### Histology

After the behavioral experiments were completed, mice were deeply anesthetized with avertin (2,2,2-tribromoethaol 1.25%, 2-methyl-2-butanol 0.78%; 20 µL/g, i.p.; Sigma Aldrich) and transcardially perfused with PBS followed by 4% paraformaldehyde in PBS. Brain were removed and postfixed overnight at 4°C. Then, 40-µm coronal slices containing the striatum were cut with a vibratome (VT1000 S; Leica) and stored in PBS. Sections were washed for 10 min in PBS, incubated for 10 min with Hoechst 33258 (1:10000 in PBS), and mounted on glass slides and coverslipped with Fluoromount. Slices were imaged using a microscope (V16; Zeiss) confirming for all mice the adequate localization of the lens in the dorsal striatum.

### Identification of behaviors

To identify the behaviors that mice displayed during open-field explorations recorded through EthoVision XT14 (Noldus), we combined deep learning tools and clustering methods to generate a predictive model for labelling behaviors. First, the x-y coordinates of different mouse body parts (nose, neck, left ear, right ear, microendoscope camera, body center, tail start, and tail end) were identified using a DeepLabCut^46^ deep neural network trained using 800 randomly selected and manually annotated frames taken from 40 different videos. The training regimen was set to the DeepLabCut default^46^. Any coordinate detected by DeepLabCut with a likelihood of less than 0.9 was removed from further analysis. For each video frame, the above body parts were used compute six features describing the mouse posture: the *body speed*, which was computed as the projection of the body center speed vector along the mouse body axis; the *head speed*, which was defined as the norm of the difference between the body center speed vector and the camera speed vector; the *movement angle*, which was calculated as the angle between the body center speed vector in the previous and subsequent frames; the *body length*, which was calculated as the sum of the distance between the neck and body center and the distance between the body center and tail start; the *neck elongation*, which was measured as the distance between the neck and body center; and the *head elevation*, which was calculated as the distance between the neck and the point defined by the orthogonal projection of the camera position along the vector orthogonal to the vector defined by ears positions. The temporal evolution of these six features was smoothed over 20 frames. Then, using a quarter of the data points, a nonlinear dimension reduction algorithm (t-distributed stochastic neighbor embedding, t-SNE) was applied to identify potent clusters in 3 dimensions. Ten replications of the t-SNE algorithm were computed for the same data. The clusters were then identified using a Gaussian mixture model (GMM). Fifty replications of the GMM clustering with random initializations were calculated for each individual t-SNE replicate. The resulting 500 classifications of the frames were subsequently clustered using the Hamming distance to identify groups of frames that were consistently classified together by the tSNE and GMM methods. Clusters with less than 20 frames were removed. The remaining clusters were merged in ascending order using the Wasserstein distance (aka earth mover’s distance) until a cutoff of 1 was reached. The resulting clusters were manually registered by visually inspecting the corresponding video frames and evaluating the distribution of cluster elements in the feature space as one of the following behaviors: *locomotion fast* (body speed greater than ∼15 cm/s); *locomotion straight* (nonzero body speed, movement angle of approximately 0); *locomotion turn right* (nonzero body speed, nonzero positive movement angle); *locomotion turn left* (nonzero body speed, nonzero negative movement angle); *still turn right* (body speed of approximately 0 cm/s, nonzero positive movement angle); *still turn left* (body speed of approximately 0 cm/sec, nonzero negative movement angle); *head up* (body speed of approximately 0 cm/s, small neck elongation, high head elevation); *rearing* (body speed of approximately 0 cm/s, small body length, small neck elongation, high head elevation); *grooming* (speed of approximately 0 cm/s, nonzero camera speed, small body length, large movement angle variations); *locomotion sniffing* (nonzero body speed, large body length, high neck elongation); *still sniffing* (body speed of approximately 0 cm/s, large body length, high neck elongation); or *immobility* (body speed of approximately 0 cm/s, camera speed approximately of 0 cm/s). Finally, using these defined clusters and their distributions in the feature space, we evaluated for each frame its likelihood of belonging to each behavior cluster. The behavior was determined according to the highest likelihood. Any behavior episode of less than 100 ms (i.e., 4 frames) was removed. In cases in which some video frames were unlabeled, the first half of these series were attributed to the previous behavior, while the second half was attributed to the following behavior. The resulting identification of spontaneous mouse behavior in the open field exploration was systematically visually inspected to ensure proper classification.

### Calcium signal extraction, deconvolution and longitudinal cell registration

The calcium movies were preprocessed for spatial binning (downsampled by 4; OpenCV, Python) and subsequently motion-corrected and analyzed using CaImAn^18^ to take advantage of the capabilities offered by the constrained nonnegative matrix factorization for endoscopic data (CNMF-E) algorithm to estimate and correct for background neuron somata^19^. The temporal CNMF-E components were manually curated to remove components with poor signal-to-noise ratios (peak-to-noise ratio of less than ∼3), large baseline fluctuations, or inappropriate spatial footprints. The selected temporal calcium components were also deconvolved to estimate spike trains in the calcium measurements using MLspike^20^. To register cells across imaging sessions for the same animal, we used CellReg^29^, which uses a probabilistic method to automatically register cells that are present in two or more recordings from the same mouse.

To control for longitudinal registration, two control lists of registered pairs of cells between pairs of sessions were generated: the first relied on a random shuffling of registered neurons in one of the two sessions, while the second relied on replacing all the neurons identified in one of the two sessions with their closest neighbor.

### Quantification of neuronal activation similarity and behavioral similarity

Two measures of neuronal similarity were used: the first compared the neuronal activation during each behavior over time during one recording session (or between recording sessions using pairs of longitudinally registered cells present in both sessions), while the second compared neuronal activation between different behaviors within a given recording session. For the measure of the neuronal activation similarity for each behavior during one session (30 min long), we first split this session into two 15-min halves. For each time period, we calculated for each behavior the average value over time (15 min) of the deconvolved activity for each neuron, denoted as ***X****_1_* and ***X****_2_*. The similarity was evaluated as 1 – || ***X****_1_*/||***X****_1_*|| – ***X****_2_*/||***X****_2_*|| ||, where ||***X***|| is the Euclidean norm of ***X***. As a control, the same similarity metric was computed by calculating ***X****_1_* and ***X****_2_* based on all possible partitions of 30 min into two periods using 5-min-long segments. We also calculated the neuronal similarity between odd and even frames by computing ***X****_1_* and ***X****_2_* in odd and even calcium frames. Finally, the neuronal activation similarity was also evaluated for each behavior by calculating ***X****_1_* in one episode every two episodes and calculating ***X****_2_* in the remaining episodes of each behavior. The spatial shuffle similarity was calculated as the mean of 10 random permutations of indices from ***X****_2_*. In addition, the neuronal activation similarity between pairs of behaviors was calculated using the same formula as above. The average neuronal activity for each behavior was calculated over the duration of the entire session (30 min) except otherwise mentioned, and the similarity was computed for each pair of behaviors. The distance between behaviors (behavioral distance) was estimated for each pair of identified behaviors as the summation of the Wasserstein distances for each of the six features describing the mouse posture (body speed, head speed, movement angle, body length, neck elongation, and head elevation). The similarity between behaviors (behavioral similarity) was calculated as the opposite of the behavioral distance. Similar results were obtained using similarity metrics based on the Wasserstein distance or Bhattacharyya distance. To evaluate the relationship between the neuronal activation similarity and the behavioral similarity, we used the Spearman correlation coefficient.

For all of the above experiments, if the sampling duration over which the average activity was computed was less than 5 s, the data were excluded from further analyses.

### Support vector machine decoding of behavior based on SPNs activity

To decode behaviors based on neuronal activity, we trained a set of binary support vector machine (SVM) classifiers for multiclass classification using the one-vs.-one strategy (scikit-learn Python package)^47^. The training was performed using 5-fold cross validation to predict the detected behavior time series, with the deconvolved calcium activity convolved over a 500 ms square window. The outputs of the classifiers were combined, and the behavior with the highest number of votes was identified as the most likely behavior. The decoding accuracy was estimated as the fraction of time bins during which the predicted behavior corresponded to the observed behavior. The behavior reconstruction error was calculated using the behavior distance between the predicted and observed behaviors. Alternatively, to estimate the capability of SVM classifiers to separate one given behavior from any other behavior, which is referred in the text as one behavior vs. rest decoding, we calculated the accuracy as the simple matching coefficient between the observed and predicted behaviors (i.e., the true positive prediction for this behavior and the true negative prediction for this behavior).

The chance level for the decoding performance was obtained by training SVM classifiers on time-lagged data. Briefly, we flipped the behavior time series (the first element becomes the last and vice versa) and applied a cyclic permutation with a random time lag. This procedure destroys the relationships between the behavior and calcium activity time series but preserves the time correlations of the neural activity time series. For each recording session, 10 random time lags were used. For each random time lag, we trained a new set of SVM classifiers and evaluated its performance in predicting the original behavior using the original calcium time series. When plotted, the individual points for SVM decoding based on the shuffled data represent the average of the 10 random time lags.

For longitudinal predictions between pairs of recording sessions, the SVM classifiers were trained on one session using only neurons that were registered in these two sessions and the corresponding behavior time series. The decoding performance was evaluated using calcium events from the second session of cells registered in both sessions. If less than 40 neurons were identified in both sessions, the analysis was discarded.

### Detection of behavior-active neurons

To characterize the statistical significance of neuronal activation during identified behaviors, we employed a behavior information criterion that was calculated as the mutual information score between the calcium event occurrence and the mouse behavior. The behavior information for each cell was calculated using the following formula:

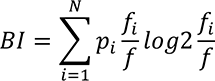

where *i* is the behavior, *p_i_* is the fraction of time spent performing behavior *i*, *f_i_* is the average event frequency during behavior *i*, and *f* is the overall average event frequency. We corrected this measure for sampling bias in the information measures by using shuffled distributions of the events. For each cell, we generated 1000 random permutations of the events and calculated a behavior information value for each permutation, thus generating 1000 random behavior information values to which the actual behavior information value was compared. We labeled a cell as a behavior-active cell if the behavior information value was more than 4 sigma from the shuffled distribution (significance of BI above 4)^25, 26^.

When this method was used in conjunction with SVM-based predictions, analyses were discarded if less than 40 neurons were labeled as behavior-active in a given session.

### Detection of behavior-silent (and behavior-inactive) neurons

To characterize cells that were consistently silent during a given behavior, we calculated for each neuron and each behavior the average event rate for all episodes of this behavior. We then evaluated for each cell and each behavior an activation occurrence value, which described how often this neuron was active during episodes of this behavior (event rate > 0). For each behavior, a neuron was labeled as behavior-silent if its activation occurrence was less than 0.025. Alternatively, for each behavior, we labeled a cell as behavior-inactive if its average event rate during this behavior was less than 0.1 Hz.

When this method was used in conjunction with SVM-based predictions, analyses were discarded if less than 20 neurons were labeled as behavior-silent (or behavior-inactive) in a given session.

### Quantification and statistical report

Unless otherwise stated, the mean ± standard error of the mean (SEM) was used to report data. For all statistics, we used a linear mixed-effects model followed by analysis of variance to account for between-subject and within-subject effects in the case of incomplete design (exclusion criteria mentioned in the above sections). To compare the dSPN and iSPN groups, post hoc analyses were performed using permutation-based t tests. For hypothesis testing, the significance was set to 0.05. Statistical analyses were performed in MATLAB (MathWorks). Animals were excluded prior to data acquisition if the imaging quality or focal plane were poor or after acquisition but before secondary analyses if movement artifacts were impossible to correct using CaImAn. The details of the statistical procedures and results are provided in Supplementary Table 1.

## Notes

### Competing Interest Statement

The authors have declared no competing interest.

